# Allele Frequencies at Recessive Disease Genes are Mainly Determined by Pleiotropic Effects in Heterozygotes

**DOI:** 10.64898/2025.12.05.692665

**Authors:** Jonathan Judd, Jeffrey P. Spence, Nikhil Milind, Linda Kachuri, John S. Witte, Jonathan K. Pritchard

## Abstract

The classic theory of mutation-selection balance predicts the equilibrium frequency of genetic variation under negative selection. The model predicts a simple relationship between the total frequency of deleterious variants, mutation rate, and strength of selection, with different functions for recessive and (co-)dominant genes. In this study, we investigate whether genes associated with human recessive disorders fit the predictions of this classic model. By comparing observed frequencies of loss of function variants (LoFs) to those expected under mutation-selection balance we find that, for nearly all recessive genes, the observed frequencies are too low to be explained by purely recessive selection. Analyzing the effects of heterozygous LoFs on quantitative traits from the UK Biobank, we find that recessive disease genes have widespread quantitative effects in heterozygotes. Together, these results suggest that most selection experienced by pathogenic mutations in recessive disease genes may be due to stabilizing selection in heterozygotes. We conclude that very few human genes follow the classic model of recessive mutation-selection balance.

## Introduction

A century ago, J.B.S Haldane developed the now-classic theory of “mutation-selection balance”, which predicts the equilibrium frequency of deleterious variants [1, 2]. The key insight of mutation-selection balance is that, at equilibrium, the rate at which new deleterious variants are generated in a gene by mutations must exactly offset the rate at which variants are removed by natural selection.

In the last decade, advances in whole-exome and whole-genome sequencing have enabled sequencing of hundreds of thousands of healthy people [3–6]. Consistent with the predictions of mutation-selection balance, these studies show that important genes are highly depleted of functional variants such as loss-of-function (LoF) mutations [3–5]. This observation has motivated a great deal of work on estimating the strength of selection against LoF heterozygotes (here denoted *s*_*het*_) for each gene, usually assuming additive models of selection [7–10].

However, the standard additive model does not apply to fully recessive genes. In this case, selection acts only in homozygotes; since homozygotes are rare compared to heterozygotes, the classic mutation selection-balance theory predicts that pathogenic variants should be found at much higher frequencies in recessive genes than in dominant or co-dominant genes.

It is generally believed that the recessive version of mutation-selection balance should apply to a wide variety of autosomal recessive diseases. Recessive diseases, including cystic fibrosis, sickle cell anemia, and Tay-Sachs disease [11–14], occur when patients inherit pathogenic mutations that abrogate function in both copies of a particular gene.

While significant progress has been made on estimating *s*_*het*_, estimating selection coefficients in homozygotes (denoted *s*_*hom*_) remains challenging due to the difficulty of disentangling heterozygous effects from homozgyous effects in population genetic studies, and due to the rarity of homozygotes and survivorship bias in observational studies [7, 15, 16].

Here, we take a different approach, simply asking whether the frequencies of putative pathogenic mutations in annotated recessive genes are compatible with recessive mutation-selection balance. After addressing several technical challenges we conclude that, for the vast majority of annotated recessive genes, the frequencies of pathogenic mutations are too low to be explained by a fully recessive model.

To explain this, we propose that most recessive disease genes also have meaningful phenotypic effects in heterozygotes — and that these pleiotropic effects are the main target of selection. At a mechanistic level, we can expect that heterozygous mutations would often reduce gene expression or functionality by 50%; this in turn might affect molecular phenotypes, such as chromatin accessibility, with cascading quantitative effects on organism-level traits [17–20]. Consistent with this prediction, we find that LoF mutations in genes annotated as being fully recessive frequently have significant effects on quantitative traits measured by UK Biobank. These heterozygous effects are likely too small to be recognized as clinical conditions, and yet strong enough to be subject to negative selection.

In summary, we report here that the frequencies of pathogenic mutations in nearly all annotated recessive genes are too low to be explained by Haldane’s classical model of recessive mutation-selection balance. Instead we propose that most selection at these genes is due to stabilizing selection acting on quantitative effects in heterozygotes.

## Results

Mutation-selection balance predicts the total allele frequency of all pathogenic variants for a gene, *p*_*path*_, by calculating the frequency at which the rate of new pathogenic alleles arising by mutation is exactly offset by the removal of alleles by natural selection:

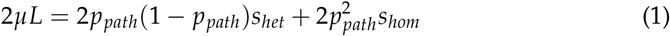

where *µ* denotes the per-base pair mutation rate, and *L* is the effective number of sites that can produce pathogenic mutations [1, 2]. When selection on heterozygotes, *s*_*het*_, is non-negligible (i.e., for dominant or co-dominant traits), nearly all selection occurs in heterozygotes and the equilibrium frequency can be approximated as

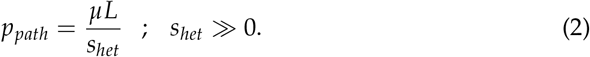

In contrast, for fully recessive genes, selection occurs only in homozygotes, so the equilibrium frequency is determined entirely by the frequency of homozygotes:

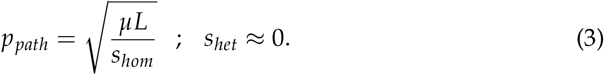

Under this model, the frequency of pathogenic alleles, *p*_*path*_, is proportional to the square root of the gene’s total mutation rate *µL* (Figure 1A). A key prediction for the present paper is that, even for fully lethal recessive mutations (*s*_*hom*_ = 1), selection cannot push the equilibrium frequency *p*_*path*_ below 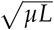 for the corresponding gene.

**Figure 1:**
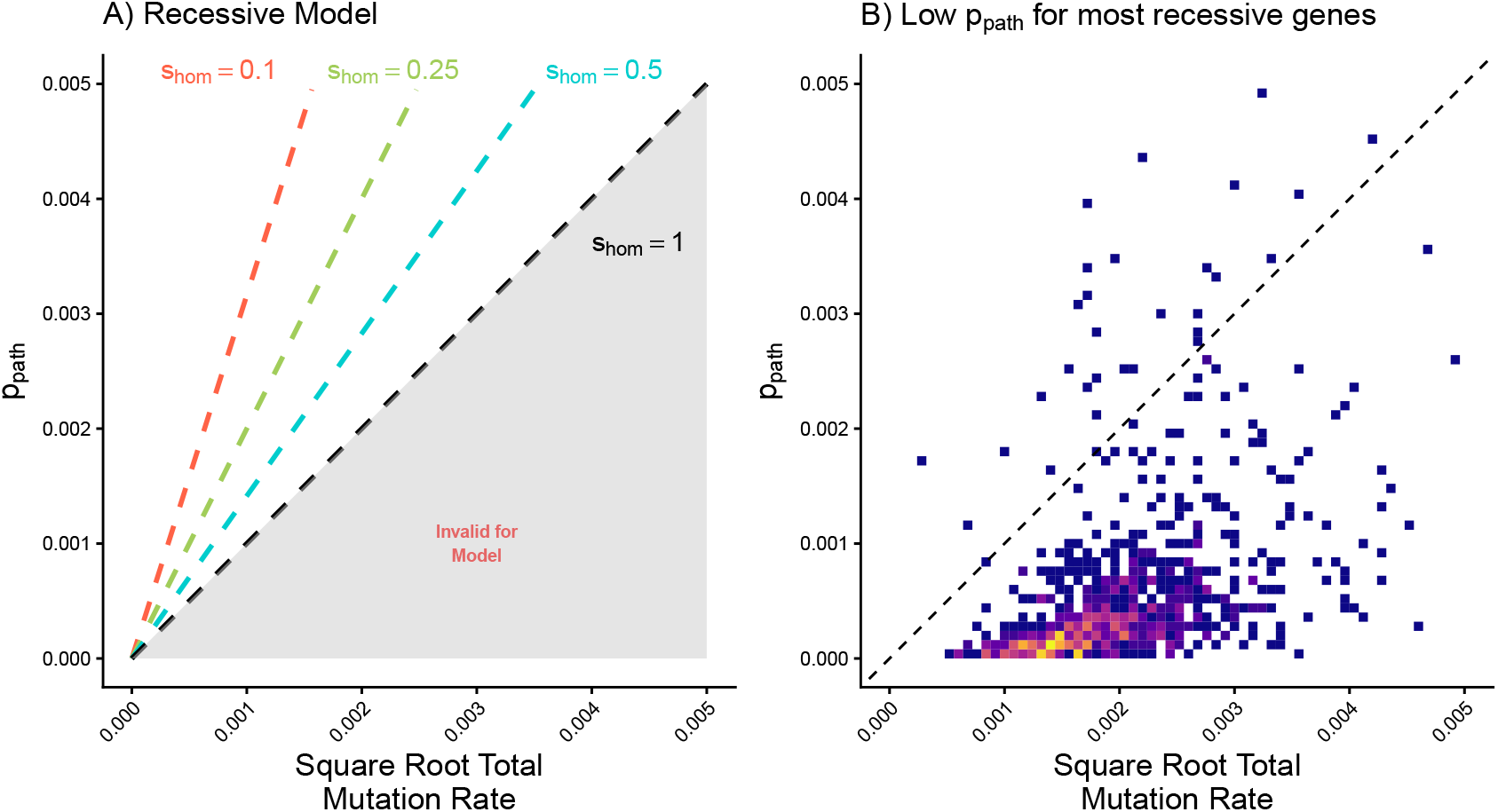
Observed Pathogenic Allele Frequency Versus Expected: **A)** Theoretical relationship between the square root mutation rate and p_path_ under recessive mutation-selection balance, for a range of values of homozygous selection, s_hom_. The shaded area represents parameter combinations that are incompatible with classic recessive mutation-selection balance. **B)** Density plot showing the observed relationship between square root total mutation rate and predicted p_path_ for recessive genes. Most genes lie in the parameter region that is inconsistent with classic mutation-selection balance.

Note that this classic derivation makes several simplifying assumptions that may not hold in practice (i.e., ignoring inbreeding, drift, and in-phase compound heterozygotes). We will consider these below.

### Curation of recessive disease genes

To assess the extent to which purely recessive mutation-selection balance pertains to human genes, we began by curating a list of genes associated with only recessive diseases. To identify the most plausible purely recessive genes, we intersected the 19,194 protein-coding genes defined from the HUGO Gene Nomenclature Committee [21], with previous lists of recessive disease genes [16, 22, 23]. We then further filtered to remove genes with non-recessive effects reported in the Online Mendelian Inheritance in Man (OMIM) database [24]. This removes 387 genes, such as *CFTR* (cystic fibrosis) and *HBB* (sickle cell anemia), that are annotated as recessive, but also have documented heterozgyous or non-recessive effects.

This results in a final list of 796 genes with the strongest support for a fully recessive mode of inheritance (Table S1).

### Comparison of *p*_*path*_ to model predictions

Previous studies of selection coefficients in heterozygotes (*s*_*het*_) have generally focused on estimating the frequencies of certain types of loss-of-function mutations that can be accurately predicted computationally, and for which it is relatively straightforward to model the corresponding mutation rates. Specifically, recent studies have focused on annotating stop-gain, splice site donor-/acceptor-disrupting variants [5, 25].

Here, we refer to these specific categories of mutations as loss-of-function (LoF) mutations, and we denote the aggregate frequency of LoF variants as *p*_*LoF*_. Gene-level information about LoF variants, including positions and allele frequencies, were retrieved from the gnomAD database v4.0, calculated separately for the Non-Finnish European (NFE), Finnish (FIN), African (AFR), and South Asian (SAS), ancestry groups [5, 25, 26]. We excluded variants with an allele frequency *>* 0.005 as previous work has shown that these are highly enriched for LoFs that are either bioinformatic errors, or may otherwise be functionally rescued [10]. (We show below that our qualitative results hold even without this filtering.)

We estimated the gene level mutation rate *µL*_*LoF*_ (to LoFs) for each gene by counting all possible LoF-causing point mutations. We then computed a context-specific mutation rate that accounts for trinucleotide context and methylation status at CpGs for each possible LoF mutation; this was averaged across all *L* possible LoF-generating mutations, to estimate a total gene-level mutation rate [5, 25].

Initial analysis showed that, for nearly all recessive disease genes, the frequency of LoFs, *p*_*LoF*_, lies far below the expected minimum frequency of 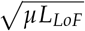 indicating that *p*_*LoF*_ cannot be predicted by mutation-selection balance under a purely recessive model (Figure S1A). However, this calculation fails to capture the fact that many individuals who are impacted by recessive disease are compound heterozygotes, carrying different mutations on each of their two gene copies. Thus, strictly speaking, the theory only applies to the full set of *all* possible pathogenic mutations, *p*_*path*_.

Compared to annotating LoF mutations, it is much more difficult to confidently identify *all* pathogenic mutations for a given gene, let alone to estimate the total mutation rate that *could* generate pathogenic mutations.

As an alternative, we estimated a scaling factor that represents the fraction of all pathogenic mutations captured by LoFs. Specifically, we identified the molecular consequences of all known pathogenic variants for 25 well-studied recessive disease genes (Figure S1B) and then calculated the percentage of this variation that is captured by our annotated LoF mutations (Figure S1C, Table S2). We found that, on average, LoFs contributed around 32% of the total number of pathogenic variants. We then used the inverse of this fraction as a conversion factor to rescale *p*_*LoF*_ and the LoF mutation rate *µL*_*LoF*_ to estimate an overall *p*_*path*_ and total mutation rate *µL* for each gene.

We were intrigued to find that, for the vast majority of recessive disease genes, the estimated *p*_*path*_ is far lower than can be explained by the recessive mutation-selection balance model (Figure 1B). This result is observed when examining the relationship between *p*_*path*_ and total mutation rate in recessive disease genes in other populations (Figure S2).

When the list of recessive genes is restricted to recessive-lethal genes, which are under the strongest homozgyous selection, we also see quantitatively similar results (Figure S3A).

Additional sensitivity analyses were performed to confirm that recessive disease gene selection and variant selection criteria, which might result in underestimating *p*_*path*_, were not likely factors to explain these observations. We tested whether the data might fit if we removed the filter on unusually high-frequency LoFs (MAF *>* 0.005), but this makes little difference to the qualitative results (Figure S3B). We also wondered whether these observations might be explained if we are underestimating the ratio of LoFs to total pathogenic mutations. However, we find that we would need to increase this factor by around 104-fold to make 50% of genes fit the model and need to increase this factor an implausible amount (1,465-fold) to make 90% of genes fit the model.

To better understand why recessive disease genes are largely inconsistent with recessive mutation-selection balance, we next considered two potential categories of explanations: deviations from the basic model including inbreeding and drift, and unexpected quantitative effects in heterozygotes.

### Potential model deviations

The classic mutation-selection balance model makes three important simplifications that may not hold in practice: no inbreeding, no drift, and modeling the gene essentially as a single site with high mutation rate. We consider each of these in turn.

#### Inbreeding

In our standard model, the probability that an individual is homozygous for pathogenic mutations follows Hardy-Weinberg expectations, i.e., 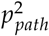. However, many populations exhibit some degree of inbreeding — i.e., relatives mate with one another more often than expected under purely random mating. Close inbreeding dramatically increases the probability that an offspring inherits a pathogenic mutation from both parents (Figure 2A). Thus, conditional on allele frequency, inbreeding increases the probability that pathogenic mutations will be exposed to selection; this results in lower equilibrium frequencies compared to random mating [27]. We wanted to assess whether this effect is strong enough to explain our results.

**Figure 2:**
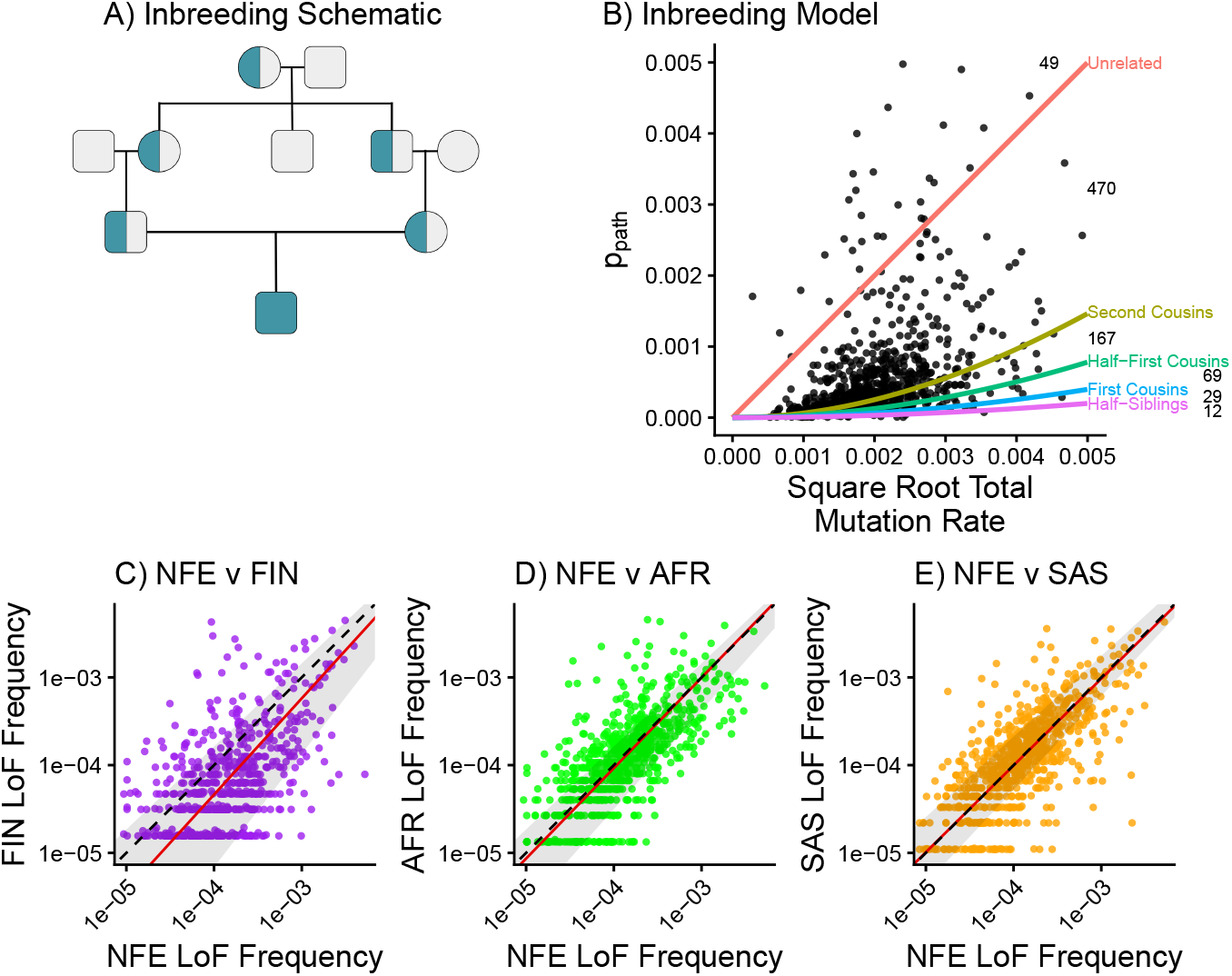
Inbreeding Under Mutation-Selection Balance: **A)** Illustration of how inbreeding, in this case between first cousins, increases the frequency of homozygotes in a population (conditional on p_path_). **B)** The lines show the theoretical equilibrium p_path_ under different levels of inbreeding; these are superimposed onto the observed data (e.g., dot is a single recessive gene). The numbers indicate the number of genes that lie between each adjacent pair of theoretical lines. Comparison of p_LoF_ of recessive genes between NFE and **C)** Finnish (FIN) populations, **D)** African (AFR) populations, or **E)** South Asian (SAS) populations, respectively. Red line represents the total least squares regression of p_LoF_ between populations. Shaded region represents 95% confidence interval of regression line. The black lines indicate y = x.

To model this, we measure the extent of inbreeding using the probability that an individual inherits both of their copies of a gene identically-by-descent, which we denote *φ*. For example, under random mating *φ* = 0; if 20% of matings are between first cousins then *φ* = 0.0125. Under this model, mutation-selection balance takes the following form in the large population size limit:

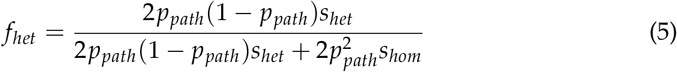

See **Appendix A** for details of the model and the derivation of the results.

Equation 4 highlights that higher inbreeding coefficients reduce the expected *p*_*path*_ for a given total mutation rate. However, the extent of inbreeding would need to be implausibly high to explain the data: to explain 95% of recessive genes, we would need to assume that *all* matings occur between first cousins (Figure 2B, Figure S4A).

For comparison, some contemporary inbreeding rates (estimated from 1000 Genomes data [28]) identify low levels of second cousin inbreeding in non-Finnish European populations and no evidence of first-cousin inbreeding [29] while others (estimated from UK Biobank Data) similarily only identify low rates of second- and first-cousin inbreeding in specific European populations [30]. This indicates that inbreeding depression is unlikely to explain the failure of recessive mutation-selection balance to fit recessive disease genes.

#### Drift in small populations

Equation 4 assumes an infinite-sized population, neglecting the effects of genetic drift on *p*_*path*_. In contrast to the additive case, genetic drift lowers the expected *p*_*path*_ under mutation-selection balance [31–34].

To address this, we derived an approximation that quantifies the effects of drift, inbreeding, and selection in homozygotes (Appendix A). Our approximation provides a good fit to forward-in-time Wright-Fisher simulations (Figure S4B). Under this adjusted model, inbreeding has a greater effect in reducing *p*_*path*_, which enables more recessive disease genes to fit mutation-selection balance for lower levels of inbreeding. However, even when accounting for drift, fitting the data for most genes would still require inbreeding on the level of ubiquitous first cousin mating (Figure S4C, Figure S4D).

We also reasoned that if either of these demographic factors (inbreeding or drift) is somehow much larger than we expect, then it would likely vary in magnitude across populations. We would not expect that the precise parameters are the same in diverse populations. But when we analyze the fit for *p*_*path*_ in diverse human populations we find only slight variation among populations and no clear bias (Figure 2C, Figure 2D, Figure 2E), a result previously described by Stolyarova *et al*. [26]. While there is evidence of a higher value of *φ* in South Asian (SAS) populations [29, 35], this does not result in the systematic reduction of *p*_*LoF*_ relative to populations with lower levels *φ* that we would expect to see under a purely recessive model. The quantitative similarities among diverse populations suggest that a biological explanation is more likely than a demographic explanation.

#### In-phase compound heterozygotes

Lastly, the standard model does not precisely model compound heterozygotes. These occur when an individual inherits a pair of pathogenic mutations at different sites within the same recessive gene. Typically, the individual would have the disease if the two mutations are out-of-phase (i.e., damaging both gene copies, but not if the mutations are are in-phase (i.e., damaging only one gene copy). Since selection would not act against in-phase compound heterozygotes, the net effect of this is to weaken selection, and *increase* the expected *p*_*path*_ [4, 36]. *Since this effect is in the opposite direction of what we see, we do not consider this scenario further*.

### Evidence for heterozygous effects

We next considered an alternative model in which pathogenic variants within recessive disease genes have effects in heterozygotes that are non-clinical, yet large enough to be relevant for selection. Since heterozygotes are so much more common than homozygotes, even weak selection in heterozygotes may be relevant for determining *p*_*path*_ [16].

An emerging body of work shows that, for most genes, the LoF frequency is determined by stabilizing selection against heterozygotes in a high-dimensional space of quantitative traits (Figure 3A). In this model, the selection against LoF heterozygotes (*s*_*het*_) is proportional to a sum of squared effect sizes of the LoF on all traits [37–41].

**Figure 3:**
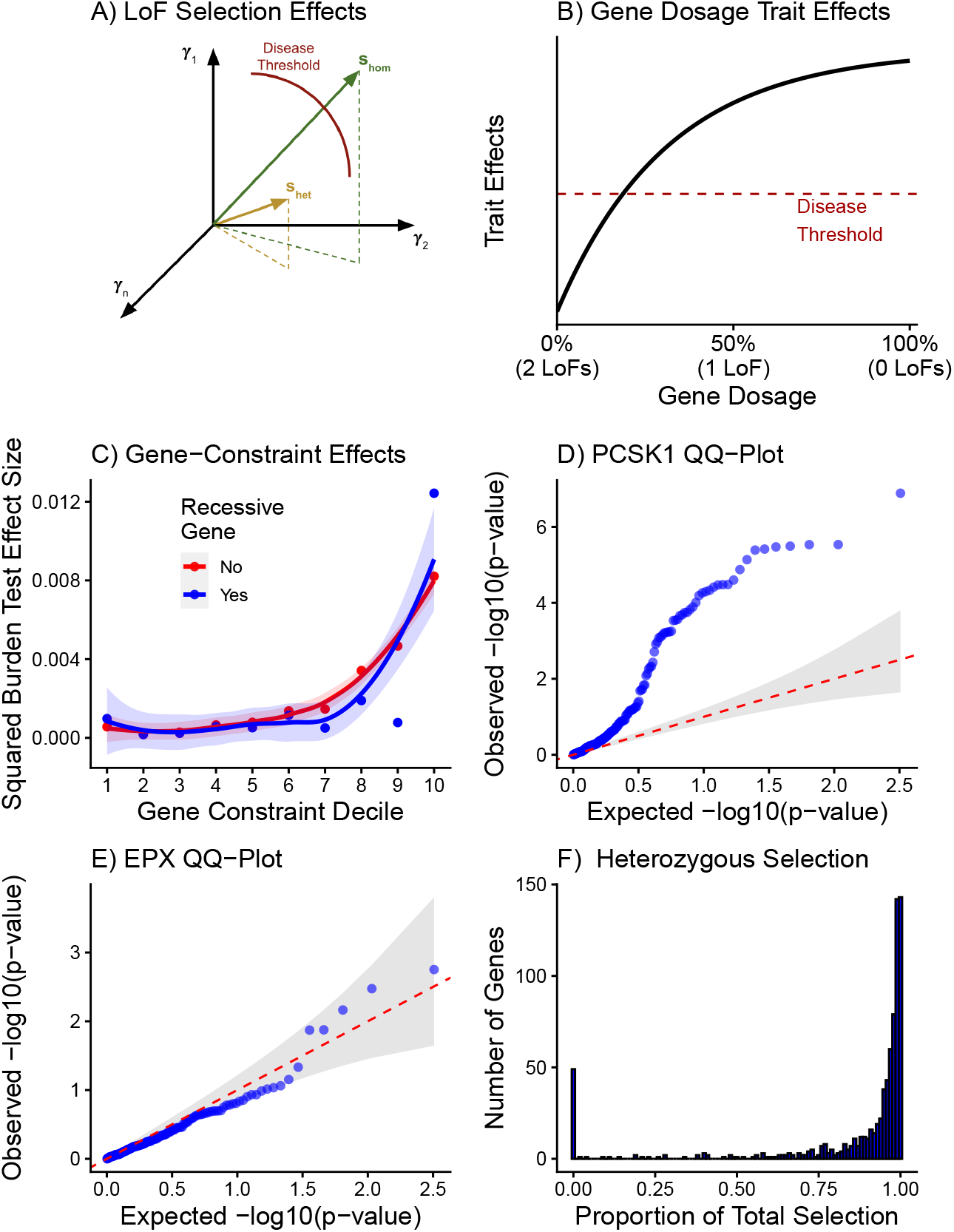
Heterozygous Burden Effects for Recessive Genes: **A)** Variant effects can be conceptualized as vectors in a high-dimensional trait space [37], shown here for hypothetical heterozygous and homozygous LoFs. Here, only the homozygous LoF crosses a clinical threshold for disease, but both genotypes are subject to selection (selection is proportional to the squared magnitude of the vector). **B)** Hypothetical relationship between gene expression dosage and model trait effects [19]. The red line represents the trait effect level where a clinical diagnosis would be made. **C)** Relationship between estimated heterozygous gene constraint decile and average bias-corrected squared LoF burden test effect sizes for 161 quantitative traits, split according to whether or not the genes are classified as recessive. Lines show loess curve fits and standard errors describing the relationship between constraint and squared burden effect sizes. **D)** Quantile-quantile plot of 161 LoF burden test p-values for the recessive PCSK1 gene. **E)** Quantile-quantile plot of 161 LoF burden test p-values for the recessive EPX gene. **F)** Histogram showing the numbers of recessive genes with a given proportion of total selection acting through heterozygous 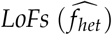.

How might this model be relevant to recessive genes? To conceptualize this, we consider the molecular effects of variants that change expression of a relevant gene between 0% (e.g., homozygous LoFs) to 50% (e.g., heterozygous LoFs) to 100% of typical expression. We model these different expression levels as corresponding to different expected values in one of the trait dimensions (Figure 3B) [18–20]. We propose that, for recessive disease genes, a single LoF may impact quantitative traits, while being insufficient to cause a clinical disease state (Figure 3B).

To test this model we sought to measure the effects of heterozygous LoFs at recessive genes on quantitative traits measured by the UK Biobank. We performed LoF burden tests on 19,194 genes, including our curated list of 796 recessive disease genes, for 161 quantitative traits. To ensure that we specifically capture the role of heterozygous LoFs on these quantitative traits, we specifically excluded homozygous LoF individuals from the data analysis.

Our group has previously shown that considering all genes there is a strong relationship between selective constraint and the average squared effect size of LoFs on quantitative traits [41]. When we analyzed the recessive genes in the same way, we find that the heterozygous squared effect sizes are extremely similar to the overall distribution across all genes (Figure 3C).

This result suggests that widespread heterozygous effects on quantitative traits in recessive disease genes in may drive the globally poor fit of the recessive mutation-selection balance model.

We also examined the gene × trait significance testing for each recessive gene on each of 161 quantitative traits. Although this test is underpowered due to low LOF frequencies and generally modest effect sizes, we find that 142 of 796 (17.8%) recessive disease genes have significant effects on one or more traits in heterozygotes (Aggregated Cauchy Association Test (ACAT) p-value < 0.05) (Figure S5A). We find qualitatively similar results when we restrict this analysis to 21 uncorrelated quantitative traits (11.8% of traits significant; Figure S5B). These percentages are similar to what we find for the heterozygous effects of all non-recessive genes using the same LoF burden test results (14.6 % for all traits; 14.0 % for uncorrelated traits) (Figure S5C, Figure S5D).

One example of a recessive disease gene with significant heterozygous effects is *PCSK1*, a gene in the same family as *PCSK9* (known for its effects on LDL cholesterol levels [42]). While *PCSK1* is known to cause endocrinopathy in an autosomal recessive fashion [43, 44], we find that it has effects on BMI, fat accumulation, and other weight-related phenotypes in heterozygotes (Figure 3D, Figure S5E, Table S3). This indicates that while homozygous LoF of *PCSK1* results in clinical endocrinopathy, variation in *PCSK1* expression such as through heterozygous LoFs have effects on quantitative traits, potentially resulting in selection against such variants [45–47].

In contrast, the recessive gene *EPX* has no observed heterozygous effects and the observed LoF frequency fits recessive mutation-selection balance. *EPX* plays a role in tyrosine nitration granule proteins in resting eosinophils and is recessively linked to eosinophil peroxidase deficiency (Figure 3E, Figure S5F) [48]. While extensive research has identified an association between *EPX* and various phenotypes, such as asthma, allergic response, and airway function, these relationships have primarily been identified in studies with total gene knockouts [48–51]. However, GWAS have identified associations between *EPX* and molecular traits, such as eosinophil and basophil count [52], suggesting that even this gene likely has some degree of phenotypic effects at small-effect variants.

Lastly, we sought to estimate, for each recessive gene, what fraction of selective removal of pathogenic mutations occurs in heterozygotes, which we refer to as *f*_*het*_, and defined:

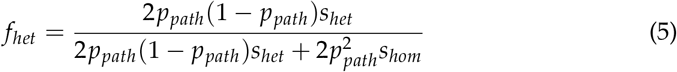

Since *f*_*het*_ is under-determined, we set

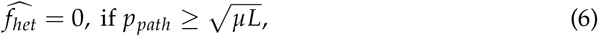

implying that the allele frequencies are consistent with fully recessive selection; and otherwise we compute a lower bound on *f*_*het*_ as follows:

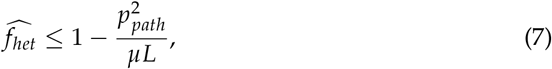

which makes the conservative assumption that homozygotes have zero fitness, i.e., *s*_*hom*_ = 1 (**Appendix A**).

Applying this to empirical data, we find that for most genes, most selection acts through heterozygotes, even when we assume LoFs are homozygous lethal. Out of all recessive genes, only 74 genes (9%) are consistent with primarily being driven by homozygous selection (Figure 3F). In contrast, we estimate that for **73**% of annotated recessive genes, at least 90% of the selection occurs in heterozygotes.

### High-frequency recessive genes

Although most recessive disease genes do not appear to follow recessive mutation-selection balance, a handful of genes in each population do have predicted pathogenic variation frequencies within the range consistent with the fully-recessive model (Figure 4A). While the specific sets of high-frequency genes vary (Figure 4B, Figure 4C, Figure 4D, Table S4), the absolute numbers are quite similar across populations (**49 in NFE, 34 in FIN, 39 in AFR, and 43 in SAS)**.

**Figure 4:**
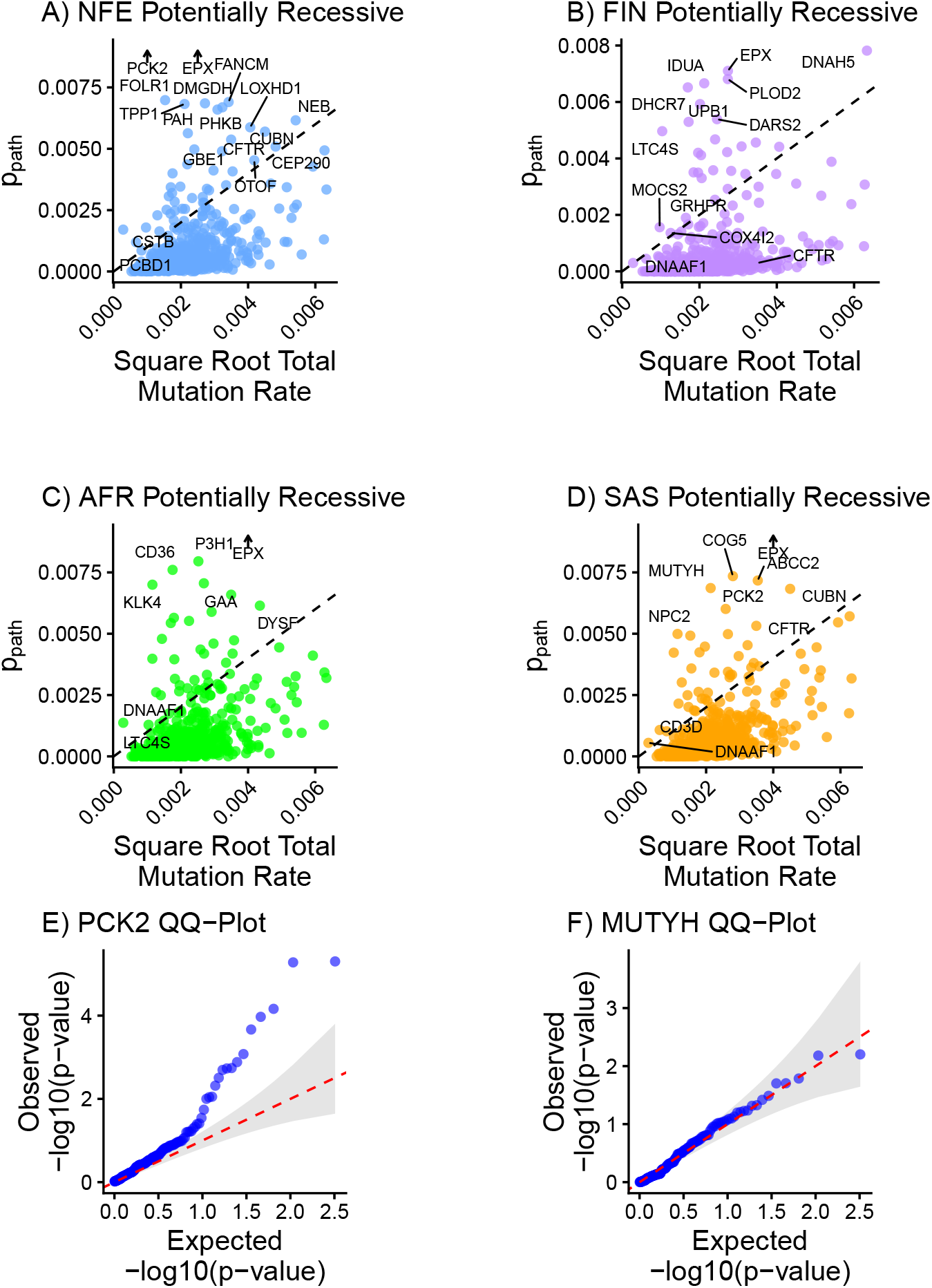
Recessive Genes Under Potentially Recessive Selection: Observed relationship between square root total mutation rate and predicted p_path_ for recessive disease genes. Chosen recessive disease genes with a predicted p_path_ in the expected range for recessive mutation-selection balance are labeled using **A)** non-Finnish European (NFE) allele frequencies, **B)** Finnish European (FIN) allele frequencies, **C)** African (AFR) population allele frequencies, and **D)** South-Asian (SAS) population allele frequencies. **E)** Quantile-quantile plot of 161 LoF burden test p-values for the recessive PCK2 gene. **F)** Quantile-quantile plot of 161 LoF burden test p-values for the recessive MUTYH gene.

Genetic drift, and potential variant ascertainment bias [26] may explain the differences between populations, and this stochasticity may result in some genes having higher than expected *p*_*path*_. Alternatively, the difference in which genes are consistent with no selection against heterozygotes may be driven by different selective pressures in different populations, for example due to differing environments. Finally, here we only considered selection *against* LoFs, but these genes with high *p*_*path*_ could also be driven by balancing selection or positive selection [53, 54].

When we examined in different populations genes previously believed to affected by balancing selection [53, 55–58], we observed variation in the genes that fit recessive mutation-selection balance (Figure S6A, Figure S6B, Figure S6C, Figure S6D). For example, two recessive disease genes that fit mutation-selection balance based on the study population and are believed to be affected by balancing selection are *PAH* (Figure S6A) and *CFTR* (Figure S6D), which are the genes associated with causing phenylketonuria (PKU) [59] and cystic fibrosis [56], respectively. If these genes and other recessive disease genes are under the influence of balancing selection, their *p*_*path*_ would be higher than expected under a model of purely recessive selection, and potentially fit recessive mutation-selection balance based on the specific population analyzed, due to differences in selective pressures, and choice of variant or LoF inclusion criteria.

Alternatively, this analysis may identify genes where the selection pressures acting on the gene and affecting *p*_*path*_ are unknown. A potential example among these recessive genes is *PCK2*, a mitochondrial isozyme of *PEPCK*, which has a well-known function in gluconeogenesis [60]. *PCK2* is consistent with having no selection against heterozygotes in both the NFE and SAS populations (Figure 4A, Figure 4D), and possesses particularly high *p*_*path*_ in the NFE population (Figure 4A, Table S4). When examining the phenotypic effects of this gene, we observe a wide range of heterozygous effects on the 161 traits we analyzed (Figure 4E), and other research has found that over-expression of *PCK2* is linked to lung cancer risk, diabetes, and obesity [61, 62]. This suggests that perhaps any selection against *PCK2* in heterozygotes caused by these phenotypic consequences is offset by some other, unknown fitness advantage.

All together, these results indicate that even these genes that are consistent with purely recessive selection may have effects in heterozygotes, and need not follow the textbook definition of recessive selection.

On the other hand, some genes appear to be purely recessive by all measures we examined. For example, *MUTYH* fits recessive mutation-selection balance (Figure 4D) and has no observed heterozygous effects (Figure 4F). *MUTYH* has a role in base excision repair and is recessively associated with colorectal adenoma risk [63]. Recent research has found that *MUTYH* continues to function in suppressing mutations even in the heterozygous LoF state [64], and only when two LoFs are present does the rate of *de novo* mutations increase, resulting in increased cancer risk. Overall, it seems plausible that *MUTYH* is an example of a gene governed by recessive mutation-selection balance.

We also examined the subset of recessive genes that match the predictions of purely recessive mutation-selection balance in the NFE, FIN, AFR, and SAS populations (Figure 4A, Figure 4B, Figure 4C, Figure 4D) to identify whether these genes showed any particular patterns. The associated diseases as well as cellular functions of these genes were varied and not limited to a specific biological function or system. However, genes that fit mutation-selection balance in the Non-Finnish European population are enriched compared to all recessive disease genes for polarized epithelial cellular component GO terms (Table S5, p-value < 0.05), which are involved in transport and environmental sensing and have associated with a variety of diseases including immunological effects [65–67], possibly indicating positive or balancing selection driven by host/pathogen competition [68–71].

## Discussion

The role of heterozygous and homozygous selection on the equilibrium allele frequency of pathogenic variation in a gene has been understood theoretically for nearly a century [2]. Purely recessive mutation-selection balance is a classic result found in textbooks and often taught in introductory population genetics courses [72, 73]. Meanwhile, the study of Mendelian autosomal recessive genes might lead one to believe that many genes should be governed by purely recessive mutation-selection balance.

Here, we find purely recessive mutation-selection balance is a poor fit for nearly every protein coding gene in the human genome, including recessive disease genes. Our analyses reveal that inbreeding is an unlikely explanation for this phenomenon, and observed *p*_*LoF*_ are instead likely shaped by selection against heterozygotes. We find that heterozygous LoFs in recessive disease genes have quantitative effects that are similar to genes overall. While these effects might not rise to the level of being classified as Mendelian disorders, the substantially larger frequency of individuals with heterozygous LoFs causes this selection against heterozygotes to be the primary determinant of *p*_*LoF*_ even in recessive disease genes.

Our results suggest that recessive disease genes are often highly pleiotropic, with effects even in heterozygotes, and hence could play roles in many complex diseases. As a result, any model of selection in humans should default to assuming that selection occurring in heterozygotes is a major factor.

Another aspect of our study worth considering is that the traits tested for heterozygous effects through burden tests are relatively limited. While these traits were chosen to represent a range of continuous non-disease, biochemistry, and anthropometric traits, they are far from exhaustive. Thus, our results provide a conservative estimate of the role of heterozygous pathogenic variants in recessive disease genes, and inclusion of more uncorrelated traits, particularly molecular traits, should only reveal an even broader range of heterozygous effects of variants in recessive genes [74–76].

Furthermore, it is difficult to determine which recessive genes with unaccounted for heterozygous effects and selection are classified as recessive because these genes are understudied or if their heterozygous effects are too small to be clinically relevant. For example, recessive genes such as *PCSK1* have strong heterozygous effects that are easily identified through LoF burden tests. While identifying these effects can inform our understanding of a wide range of quantitative traits, classifying these strong effect genes as recessive may simply be due to a lack of research. If these genes were excluded, the mutation-selection balance model may appear more valid in its current form. Inversely, there are a large number of recessive genes where we infer that the majority of selection acts through heterozygotes, but no significant gene-trait effects were identified. This can be partially attributed to the previously mentioned issue of a lack of tested traits or a lack of power, insofar as current datasets may be too small to identify individual heterozygous effects that are captured by measurements of selection, which account for the effect of variations in these genes over many generations. Identifying this second class of genes with known recessive effects and minuscule additive effects is particularly crucial for understanding human genetic variation.

In summary, we have explored the relationship between equilibrium allele frequency and natural selection on complex traits. Our results highlight the potential for widespread effects of heterozygous LoFs, even in recessive disease genes. These insights about recessive genes and natural selection will be informative for various downstream analyses, including the study of the genetic architecture of complex traits and investigations of the selective pressures underlying human genetic variation.

## Methods

### Identifying Recessive Genes

An initial list of 1,183 recessive genes was curated by previous studies [16, 22, 23] by inclusion in Online Mendelian Inheritance in Man (OMIM), a comprehensive compendium of human genes and genetic phenotypes [24], and manual curation through other resources such PubMed and Gene Reviews [77, 78]. To increase the likelihood that the list of recessive genes only affected traits through autosomal recessive inheritance, the initial list of recessive genes was cross-referenced against the OMIM database in December 2024, and any genes with non-autosomal recessive inheritance were excluded in order to produce the most updated list of 796 genes associated with recessive traits.

### Gene Data

Gene-level information about loss-of-function (LoF) variants, including allele frequencies and mutation locations, were retrieved from the gnomAD database [5]. For this study, we used gnomAD v4.0 genomic information from 705,148 unrelated individuals in the Non-Finnish European, Finnish, African, and South Asian ancestry groups. We defined LoFs as those predicted to cause stop-gain, slice-acceptor, and splice-donor effects using the Variant Effect Predictor (VEP; v85) [79]. When calculating gene-level LoF allele frequencies (*p*_*LoF*_), we summed allele frequencies for LoFs within the gene, excluding variants with an allele frequency *>* 0.005 to exclude LoFs with potential bioinformatic errors [10, 26]. (Our overall conclusions are robust to excluding this filter.) For each gene, *p*_*LoF*_ was calculated separately for each population. This variant inclusion/exclusion criteria potentially excludes many pathogenic missense and LoF variants that may lead to a recessive disease in these genes [80].However, the chosen variants are known to be highly likely to stop gene function and increases our confidence in examining pathogenic variation and its effect in recessive mutation-selection balance [5, 25, 26]. Total mutation rates for each gene were calculated by considering all possible LoF-causing point mutations and using a mutation rate that accounts for trinucleotide context and methylation status at CpGs as described in [5, 25]. Final population-specific *p*_*LoF*_s and total mutation rates along with the previously-determined recessive gene status were used to visualize additive and recessive mutation-selection balance.

### Adjusting Mutation-Selection Balance for Missing Variation

We wanted to estimate the proportion of pathogenic variants that might be excluded from the types of variants we considered above: stop-gain, splice-donor-disrupting, and splice-acceptor-disrupting point mutations. In particular, we identified the percentage of pathogenic variation in 25 well-known recessive disease genes represented by these classed of variants [81, 82]. Using ClinVar [83] to identify pathogenic and likely pathogenic short variation in recessive disease genes we analyzed genes including *HBB, CFTR, PAH, ATP7B, MUTYH, HEXA, GBA1, USH2A*, and *SMN1*, which are the genes associated with sickle cell anemia, cystic fibrosis, phenylketonuria (PKU), Wilson disease, MUTYH-associated gastric cancer, Tay-sachs disease, Gaucher disease, Usher syndrome, and spinal muscular atrophy, respectively [13, 14, 55, 63, 84–88]. After using the ClinVar-provided molecular consequences to identify LoFs and non-LoFs, we calculated the percentage of unique pathogenic variation that were predicted to cause stop gain, disrupt a splice donor, or disrupt a splice acceptor. For each gene, the inverse of these percentages was calculated as a conversion factor to adjust for potential missing non-LoF pathogenic variation, and we adjusted the visualizations of recessive mutation-selection balance by increasing *p*_*LoF*_ by the average of the calculated conversion factors and the square root total mutation rates by the square root of the average of the conversion factors.

To determine the conversion factors required so that 50% and 90% of recessive disease genes fit the recessive mutation-selection balance model, we identified the recessive disease gene whose ratio between the LoF allele frequency, *p*_*LoF*_, and square root total mutation rate, 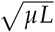, is at the 50^*th*^ and 10^*th*^ percentile of all recessive disease gene ratios, respectively. From the ratios of the genes at the 50^*th*^ and 10^*th*^ percentile, the conversion factors are the square of the inverse of the ratio for the gene (Equation 8).

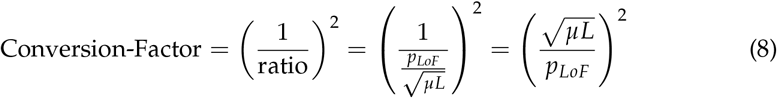

### Mutation-Selection Balance with Inbreeding

Along with the classic mutation-selection balance model (Equation 4), we adjusted another model by Nei that calculates the expected equilibrium *p*_*LoF*_ in a finite population (Appendix A) [31] in order to account for the effect of inbreeding depression on recessive genes in a more realistic population (Equation 9).

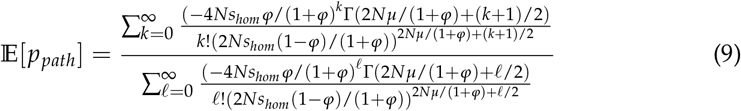

With these two models, we can predict *p*_*LoF*_ under recessive lethal selection for different total mutation rates and inbreeding values, providing a lower bound for values of *p*_*LoF*_ consistent with a purely recessive model.

### Simulating Populations with Varied Inbreeding Coefficients

In order to assess the fit of Equations 1 and 3 for varying levels of inbreeding assuming recessive lethality, we used SLiM v4.3 to generate simulated population data [89]. All simulated populations contained 20,000 individuals and possessed a single recessive lethal gene that accumulated LoF variants over 200,000 generations. We generated 125 simulated populations each with 25 varying and evenly spaced total mutation rates ([0-0.005]) and inbreeding coefficients ([0, 1/64, 1/32, 1/16, 1/8]). After simulating and calculating *p*_*LoF*_ for each population, we generated theoretical model values for the same parameters using Equation 4 and Equation 9.

The simulated data were plotted over the predicted lower bounds for LoF allele frequency under different levels of inbreeding (Equation 4 and Equation 9) to assess fit. Additionally, empirical recessive gene data was compared to both models under varying inbreeding conditions to determine the required level of inbreeding for recessive genes to be consistent with purely recessive mutation-selection balance.

### Burden Tests

We used REGENIE [90] to perform gene-level burden tests on 161 quantitative traits available in UK BioBank (UKB) [91]. For the whole-genome regression, which is the first step in REGENIE, SNPs from the genotyping array were pruned with a 1000 variant sliding window with 100 variant shifts and an *R*^2^ threshold of 0.9. SNPs were then filtered to have MAF *>* 1%, genotype missingness < 10%, and Hardy-Weinberg equilibrium (HWE) test p-value *>* 10^−15^. For LoF burden tests, we used the following covariates: age, sex, age-by-sex, age squared, 15 genotyping PCs, 20 rare variant PCs, and WES batch. After whole-genome regression, we used the second step of REGENIE to perform burden tests using the whole-exome sequencing data. We used the LOFTEE plugin in Ensembl’s Variant Effect Predictor to annotate high-confidence LoF sites [5, 79]. We further filtered any annotated LoF site with MAF <1% using a variable-frequency filter across genes based on their constraint, as described previously [10, 20]. For each gene, we filtered to individuals that carried at most one LoF variant, ensuring that burden tests were performed using only LoF heterozygotes. Phenotypes were rank-inverse-normal transformed in both the first and second step. The same covariates were used in both steps.

To generate a list of genetically uncorrelated traits, we used previously released estimates of genetic correlation based on common variants from the UKB (http://www.nealelab.is/uk-biobank/). Any traits with genetic correlation greater than 0.75 were clustered together, and traits with the most signal based on bias-corrected average squared burden effect size were picked from the clusters to generate a list of 21 uncorrelated traits.

### Measuring Burden Effects Across Gene Constraint

The estimated heterozygous gene constraint was calculated for all genes to determine if there was a positive relationship between heterozygous selection and heterozygous trait effects. For all genes, including recessive genes, the heterozygous gene constraint was estimated by dividing the gene’s total mutation rate by *p*_*Lof*_ as modeled by additive mutation-selection balance (Figure 1A). In addition, in order to account for statistical estimation bias, we calculated the bias-corrected squared effect size for all gene-trait LoF burden tests by subtracting the squared standard error of the test from the squared estimated effect size. Bias-corrected squared effect sizes were then averaged by gene to provide all genes an estimated gene constraint and average squared burden test effect size. All genes were then segregated into deciles based on estimated gene constraint. Average squared burden test effect sizes for each gene were averaged again by bin and recessive gene status. Final deciles and decile effect sizes were lastly separated by recessive gene status and plotted to visualize the relationship between constraint and heterozygous effect size.

### Calculating Proportionate Heterozygous Selection

In order to estimate a lower bound of the effect of heterozygous selection on recessive genes, we simplified the mutation-selection balance model 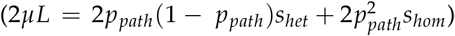 to compute *s*_*het*_ given the gene’s *p*_*path*_ and total mutation rate, and assuming the recessive gene was recessive lethal, *s*_*hom*_ = 1. Details on the estimator are provided in **Appendix A**. We then used this estimate to calculate the proportion of heterozygous selection versus total selection (the combination of heterozygous and homozygous selection), for each gene with the same assumption of recessive lethality. This provides an analytical lower bound for the proportion of total selection acting on each gene through heterozygotes.

### Recessive Gene QQ Plots for Quantitative Traits

Quantile-quantile plots were generated for all recessive genes using −log10(p-values) generated from LoF burden tests for a list of 161 UKB quantitative traits and subset of 21 uncorrelated quantitative traits. Observed −log10(p-values) were compared to expected values from a uniform distribution. 95% confidence intervals were generated from beta distribution using the 2.5% and 97.5% for the i-th smallest expected p-value of the uniform distribution.

### Aggregated Cauchy Association Tests

For each recessive disease gene, Aggregated Cauchy Association Test (ACAT) was used to combine all LoF burden test p-values for the gene [92]. We performed this analysis on different sets of burden test results: 1) combining LoF burden tests for all 161 UKB quantitative traits, and 2) combining LoF burden tests for 21 uncorrelated quantitative traits. This analysis would identify recessive genes with significant non-recessive effects using the available trait data. This analysis was repeated for all 18,398 non-recessive genes independent of the our curated recessive disease gene list. With all the non-recessive genes, ACAT was used to combine all LoF burden test p-values for each gene across the LoF burden tests for all 161 UKB quantitative traits and LoF burden tests for 21 uncorrelated quantitative traits.

### Identifying and Analyzing Purely Recessive Genes

Recessive genes that possessed a *p*_*LoF*_ greater than their respective square root total mutation rate were identified to follow classic recessive mutation-selection balance and potentially only be affected by homozygous selection. As population-specific *p*_*LoF*_ were calculated, the list of “purely” recessive genes varied by population and were individually identified.

Gene Ontology (GO) enrichment analysis was performed on recessive genes identified as “purely” recessive in the non-Finnish European, Finnish, African, and South Asian populations in order to identify shared biological, molecular, or cellular function among identified recessive genes [93, 94].

### Mutation-Selection Balance with Balancing Selection

In order to understand the potential role of balancing selection on the mutation-selection balance for genes associated with recessive diseases, we visualized the relationship between *p*_*LoF*_ and square root total mutation rate for genes previously suspected to be affected by balancing selection and are associated with only recessive diseases [53, 55–58]. We performed this analysis using observed *p*_*LoF*_ from NFE, FIN, AFR, and SAS populations available from gnomAD v4.0 genomic information.

## Acknowledgments

This work was supported by the National Institutes of Health grants R01HG014005 and R01HG008140 (to J.K.P.). UK Biobank data are publicly available by request from https://www.ukbiobank.ac.uk. The research was conducted with approved access to UK Biobank data under application number 52374 (PI: pritchard). We thank Josh Schraiber for helpful discussions, and Molly Przeworski for helpful comments on the manuscript. Anastasia Stolyarova kindly provided processed data from another study [26]. Thanks to the Pritchard Lab for many valuable comments and discussions.

## Supplemental Information

### List of Supplemental Figures

**Figure S1:**
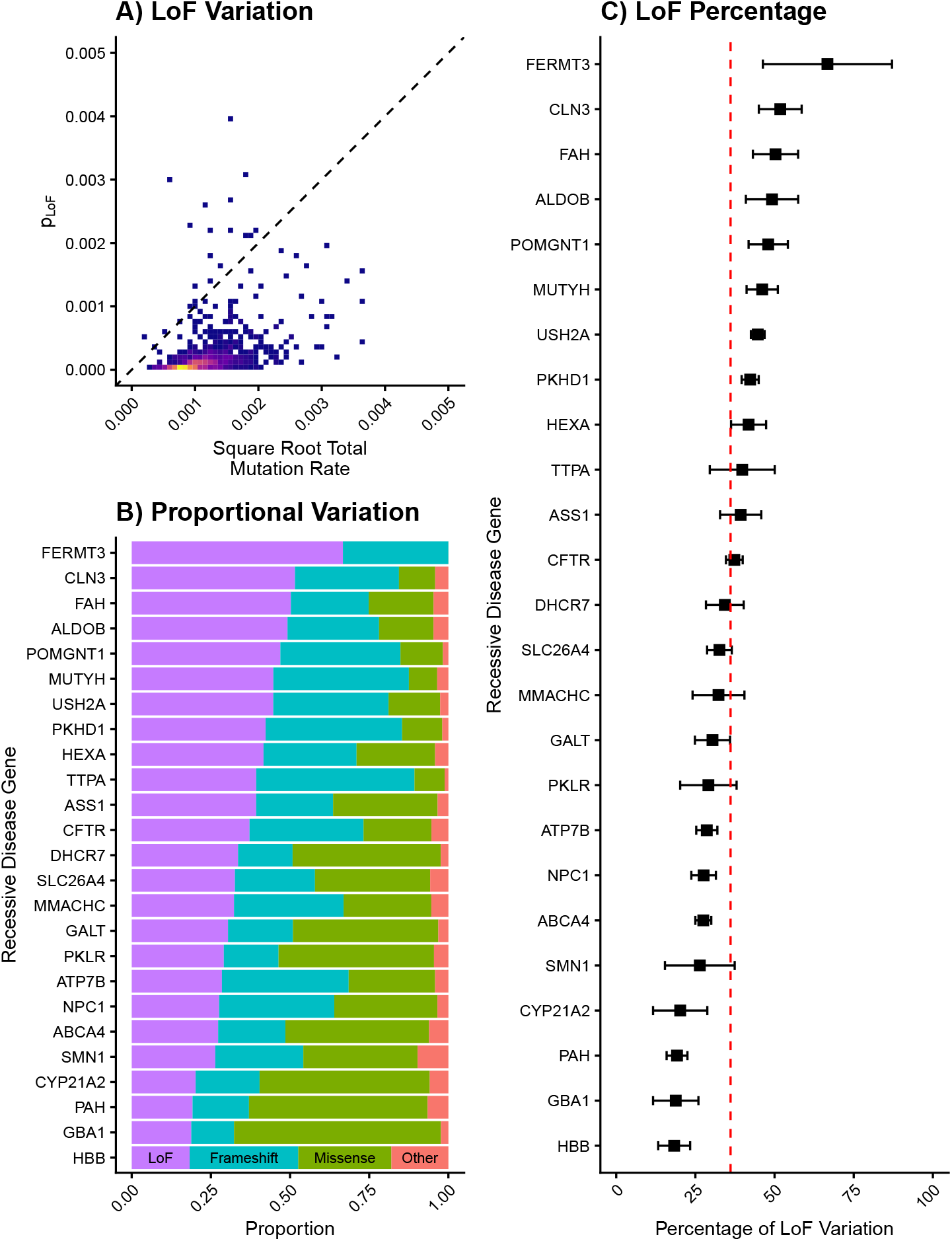
LoF and Pathogenic Variation: **A)** Density plot for observed relationship between square root mutational target size and *p*_*LoF*_ for recessive genes using NFE allele frequencies. **B)** Proportional split of observed pathogenic variation for 25 commonly-studied recessive disease genes based on predicted molecular consequence (LoF, Frameshift, Missense, or Other). **C)** Percentage of observed pathogenic variation for 25 commonly-studied recessive disease genes predicted to be LoFs. Bands represent two standard error confidence intervals surrounding LoF percentage point estimate. Red line represents average percentage of pathogenic variation predicted to be LoFs between all recessive disease genes.

**Figure S2:**
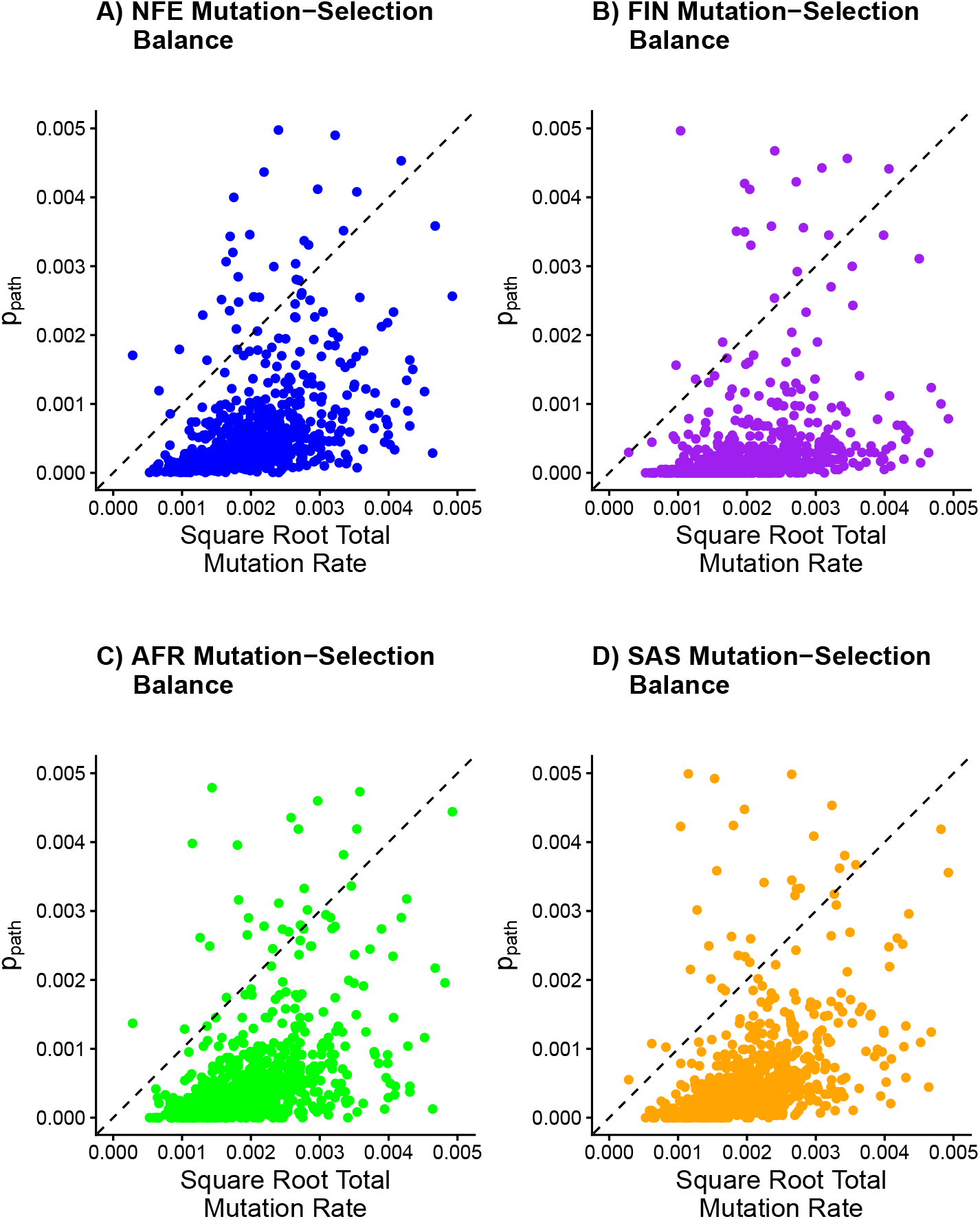
Other Population Mutation-Selection Balance: **A)** Observed relationship between square root mutational target and *p*_*path*_ for recessive genes using NFE allele frequencies. **B)** Observed relationship between square root mutational target and *p*_*path*_ for recessive genes using FIN allele frequencies. **C)** Observed relationship between square root mutational target and *p*_*path*_ for recessive genes using AFR allele frequencies. **D)** Observed relationship between square root mutational target and *p*_*path*_ for recessive genes using SAS allele frequencies.

**Figure S3:**
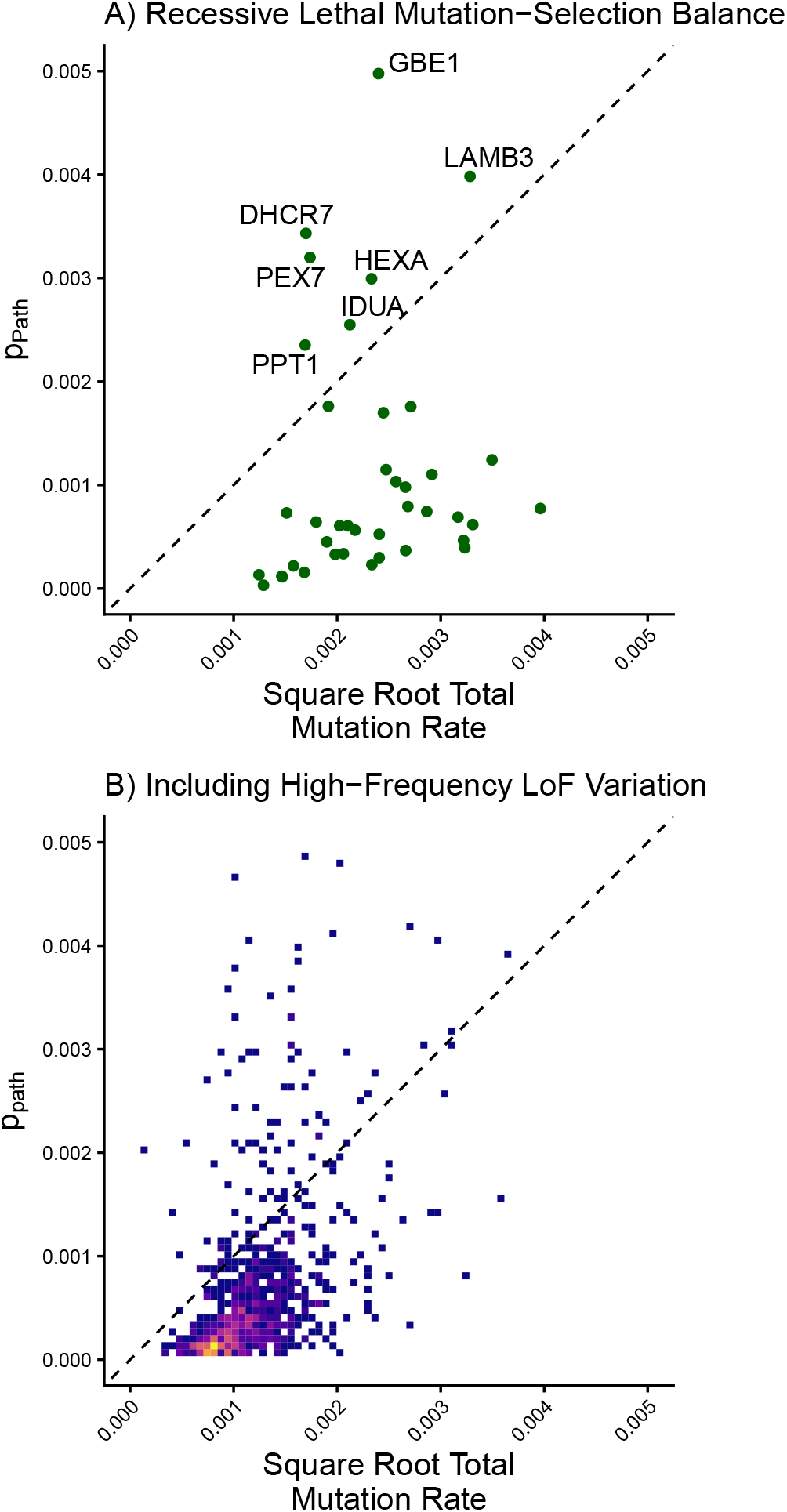
Mutation Selection Balance for Recessive Lethal Genes: **A)** Observed relationship between square root mutational target size and *p*_*path*_ for recessive lethal disease genes. Genes with a predicted *p*_*path*_ in the expected range for recessive mutation-selection balance are labeled using non-Finnish European (NFE) allele frequencies. **B)** Observed relationship between square root mutation target size and *p*_*path*_ for recessive genes using NFE allele frequencies and not excluding LoF variants with a frequency greater than 0.005 prior to calculating *p*_*path*_.

**Figure S4:**
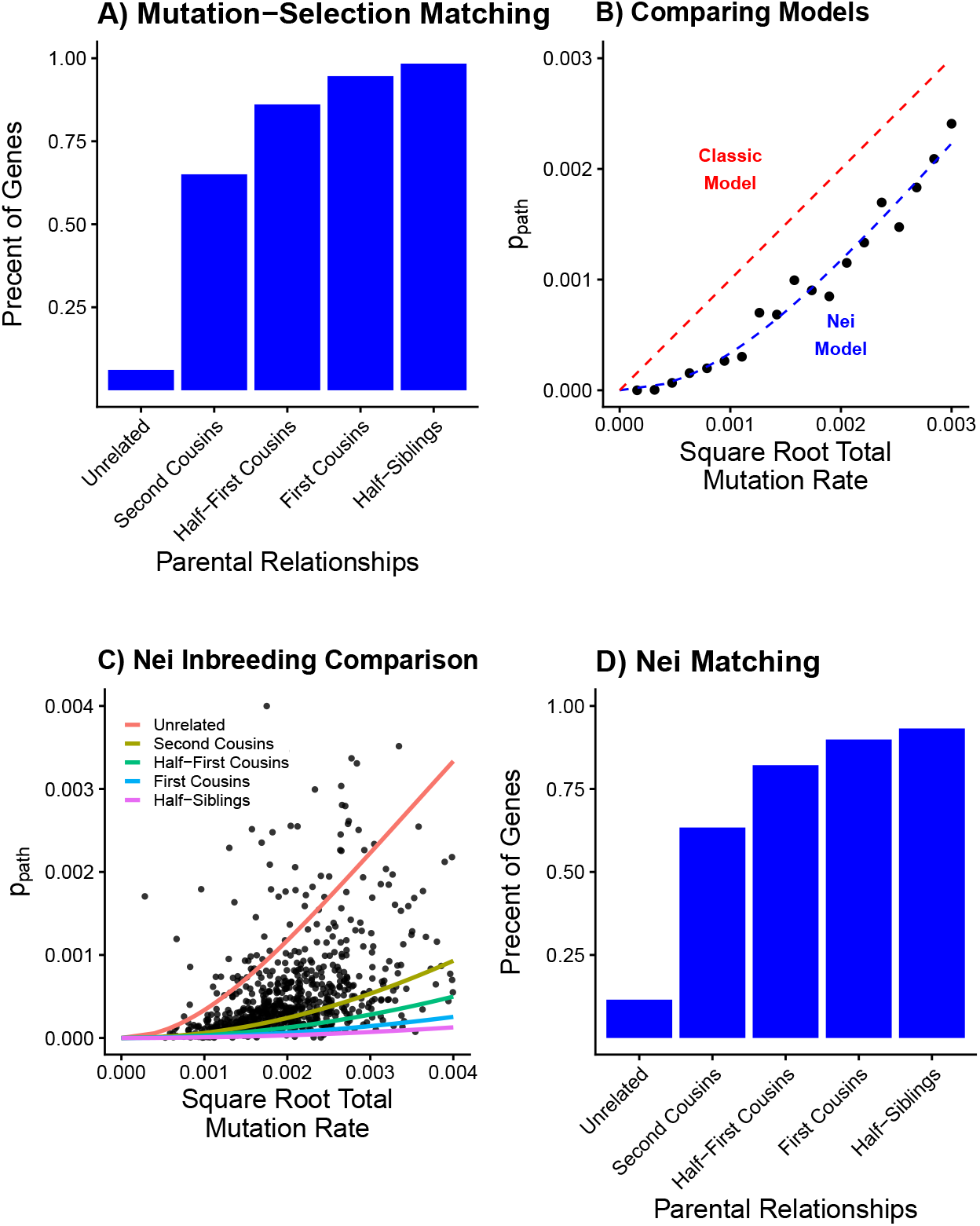
Inbreeding Models: **A)** Percentage of recessive disease genes that fit the classic recessive mutation-selection balance for different levels of population inbreeding. **B)** Comparison of the classic recessive mutation-selection balance model and adjusted Nei model (Equation 9) for a finite population for populations with no inbreeding. Lines represent predicted *p*_*path*_ for a given mutational target size for each model assuming recessive lethality. Dots represent simulated *p*_*path*_ data for recessive lethal genes with varying mutational target sizes. **C)** Observed relationship between square root mutational target size and *p*_*path*_ for recessive genes using NFE allele frequencies. Lines represent the expected relationship between square root mutational target size and *p*_*path*_ under finite population model for different levels of inbreeding in the population. **D)** Percentage of recessive disease genes that fit the finite population model for different levels of population inbreeding.

**Figure S5:**
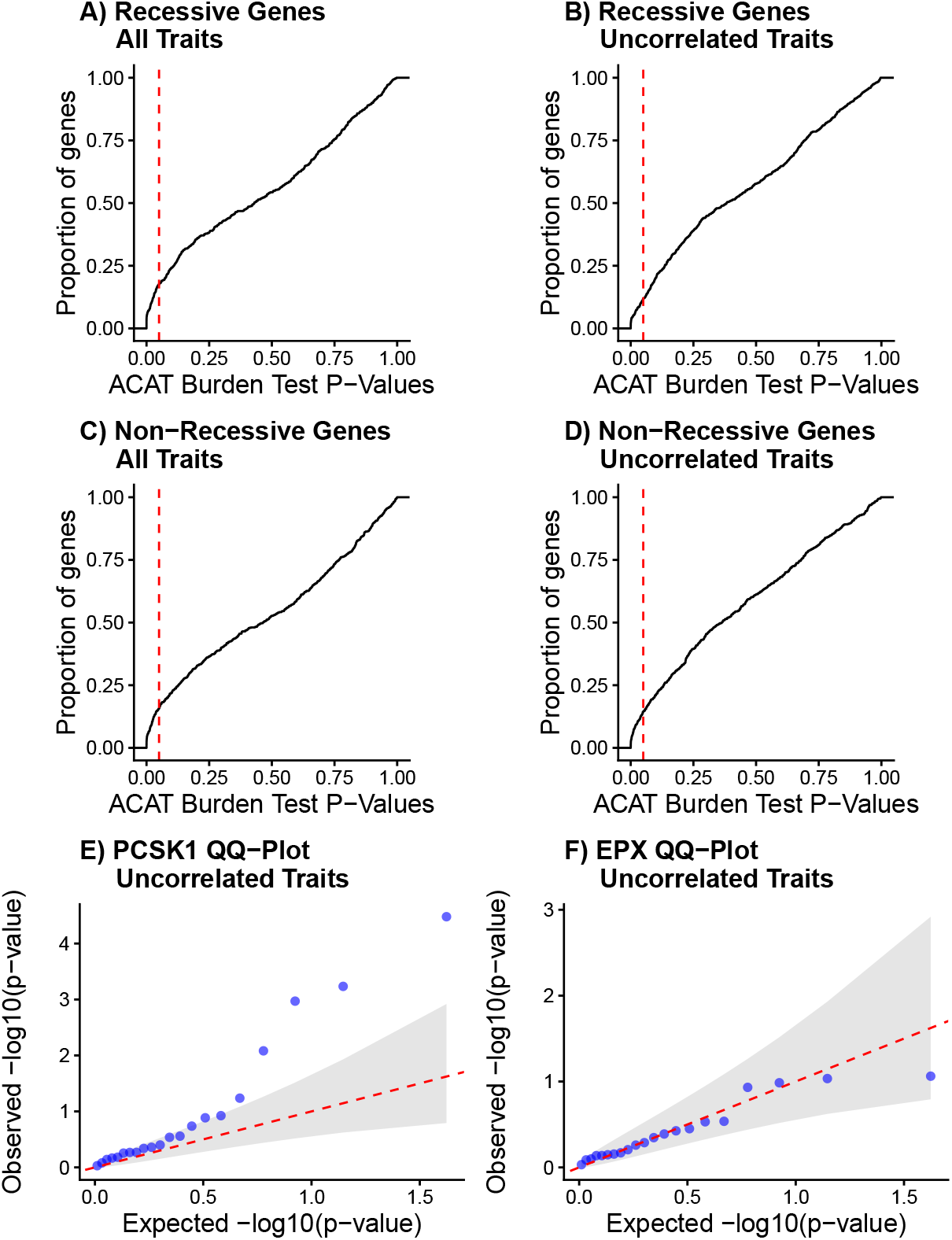
Heterozygous Effects: **A)** Empirical cumulative distribution function of ACAT omnibus test of p-values from LoF burden tests for 161 UK BioBank (UKB) quantitative traits across 796 recessive disease genes. The vertical line represents p-values that surpass 0.05. **B)** Empirical cumulative distribution function of ACAT omnibus test of p-values from LoF burden tests for 21 uncorrelated UKB quantitative traits across 796 recessive disease genes. The vertical line represents p-values that surpass 0.05. **C)** Empirical cumulative distribution function of ACAT omnibus test of p-values from LoF burden tests for 161 UK BioBank (UKB) quantitative traits across a random sample of 796 non-recessive disease genes. The vertical line represents p-values that surpass 0.05. **D)** Empirical cumulative distribution function of ACAT omnibus test of p-values from LoF burden tests for 21 uncorrelated UKB quantitative traits across a random sample of 796 non-recessive disease genes. The vertical line represents p-values that surpass 0.05. **E)** Quantile-quantile plot of 21 LoF burden test p-values from uncorrelated traits for the recessive *PCSK1* gene. **F)** Quantile-quantile plot of 21 LoF burden test p-values from uncorrelated traits for the recessive *EPX* gene.

**Figure S6:**
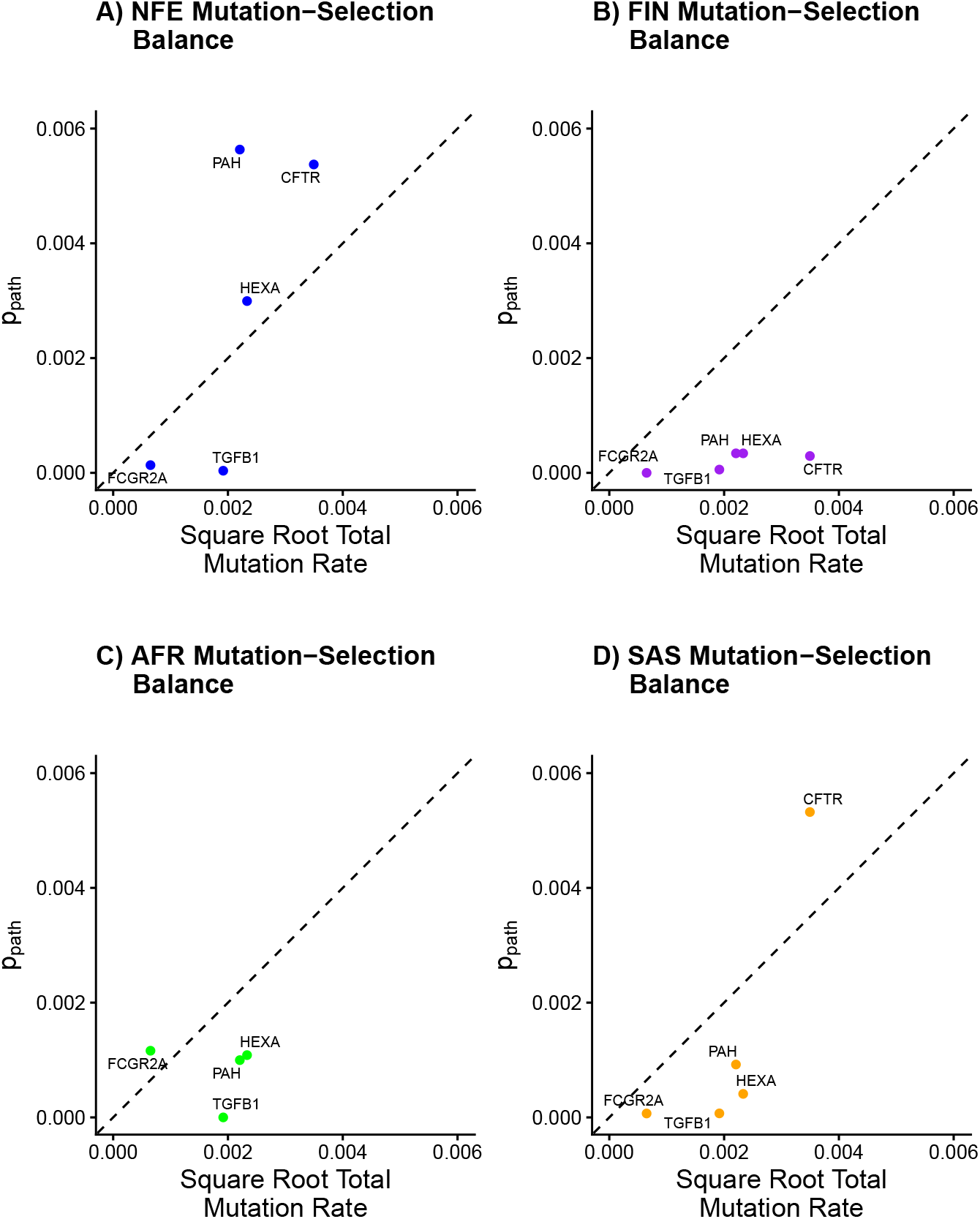
Mutation-Selection Balance Under Balancing Selection: **A)** Observed relationship between square root mutational target and *p*_*path*_ for recessive disease genes previously thought to be affected by balancing selection using NFE allele frequencies. **B)** Observed relationship between square root mutational target and *p*_*path*_ for recessive disease genes previously thought to be affected by balancing selection using FIN allele frequencies. **C)** Observed relationship between square root mutational target and *p*_*path*_ for recessive disease genes previously thought to be affected by balancing selection using AFR allele frequencies. **D)** Observed relationship between square root mutational target and *p*_*path*_ for recessive disease genes previously thought to be affected by balancing selection using SAS allele frequencies.

### List of Supplemental Tables

**Table S1: Recessive Disease Genes:** List of 796 genes associated with only recessive diseases and no other documented inheritance method listed (in OMIM as of December 2024) (Full Table in Separate File)

**Table S2:**
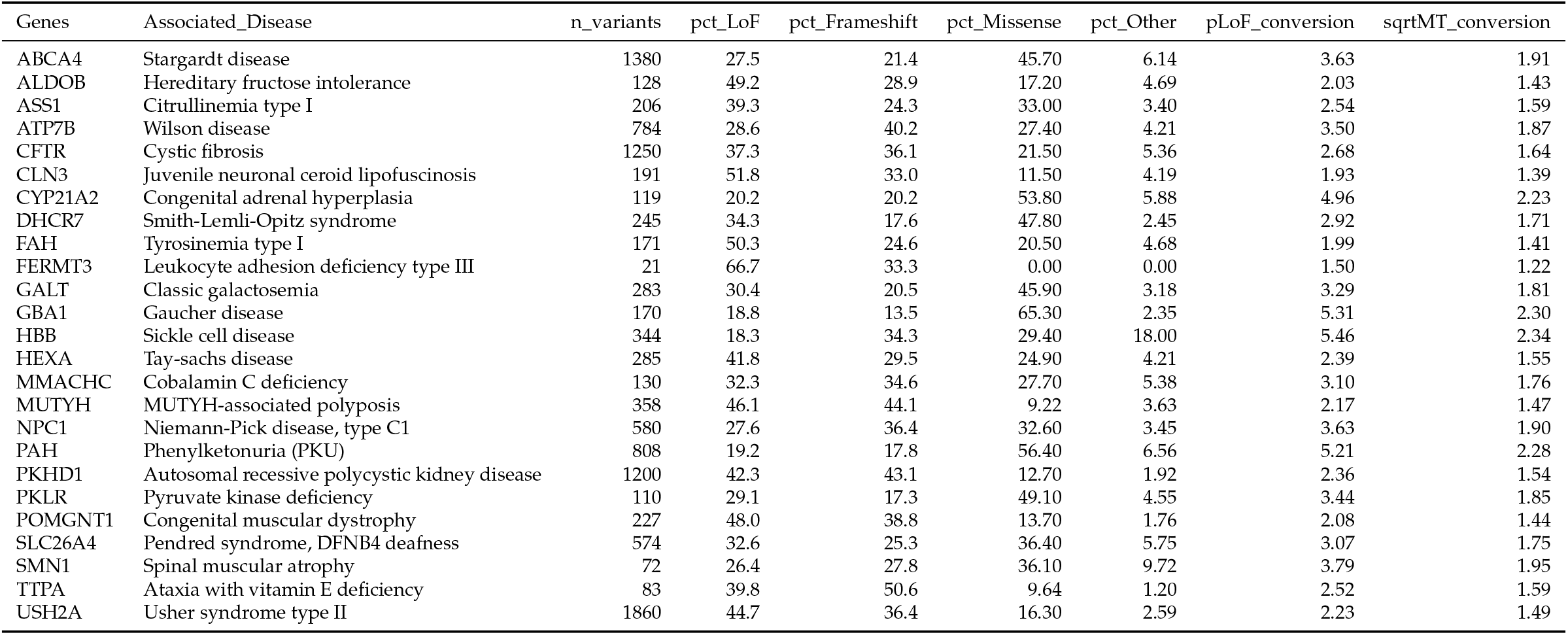
Classification of Pathogenic/Likely Pathogenic Variation for Studied Recessive Disease Genes: List of 25 commonly studied recessive diseases and genes associated with those diseases. For each gene, the number of pathogenic/likely pathogenic variants (identified through ClinVar) is listed as well as the percentage of this variation that is predicted to be loss-of-function (LoF) (nonsense, splice-acceptor disrupting, or splice-donor discrupting), frameshift, missense, or other. For each gene, a conversion factor to account for missing non-LoF pathogenic variation is included and calculated as the inverse and square root inverse of the percentage of LoF variation, respectively. Other variation defined as UTR variants, genic upstream/downstream transcript variants, inframe indels, intron variants, non-coding transcript variants, and synonymous variants

**Table S3: PCSK1 LoF Burden Tests:** PCKS1 LoF burden test results for 161 non-disease quantitative traits from the UK Biobank. Each row includes effect size estimate, standard error, and p-value for association between PCSK1 LoFs and phenotype (Full Table in Separate File)

**Table S4:**
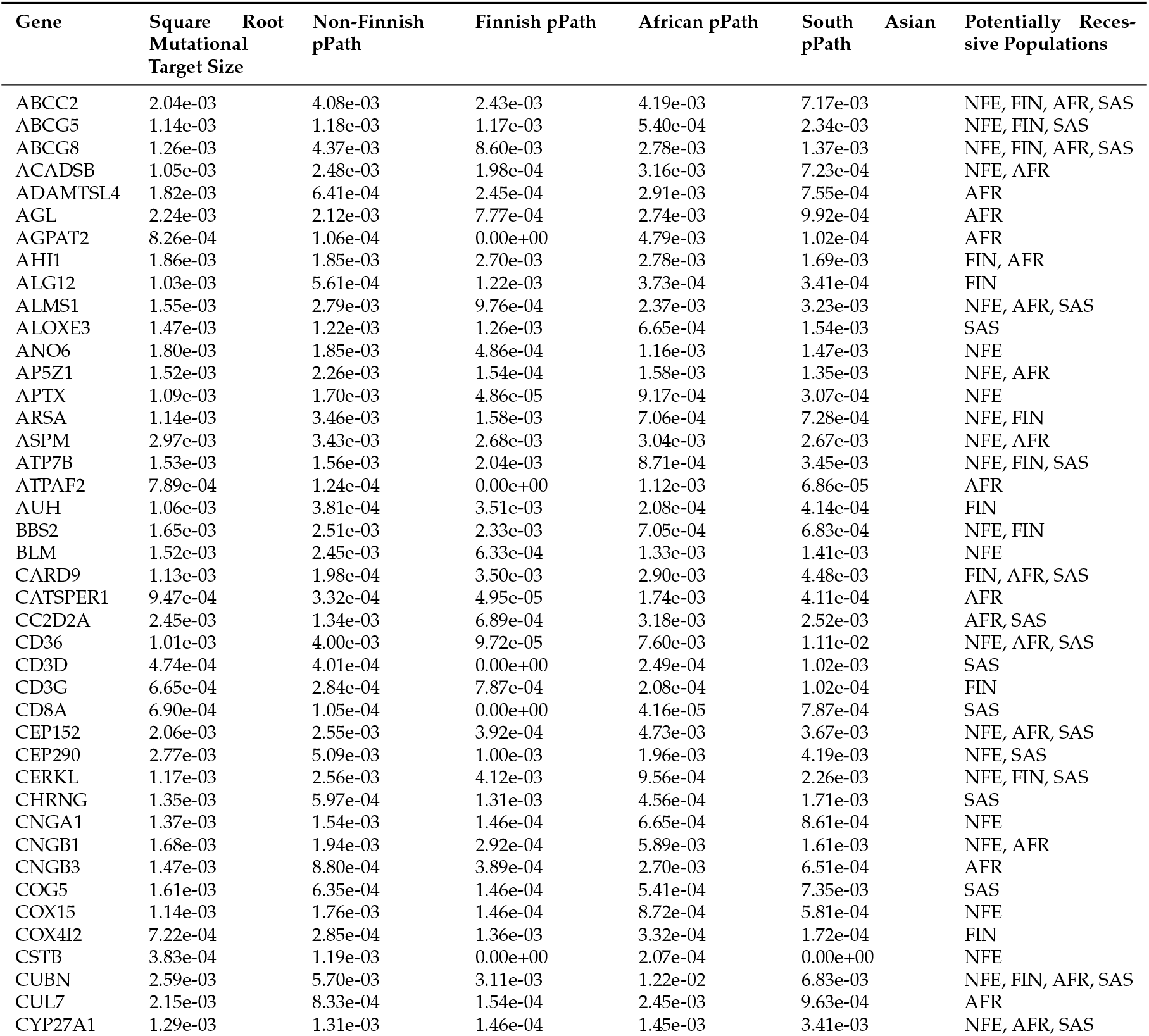
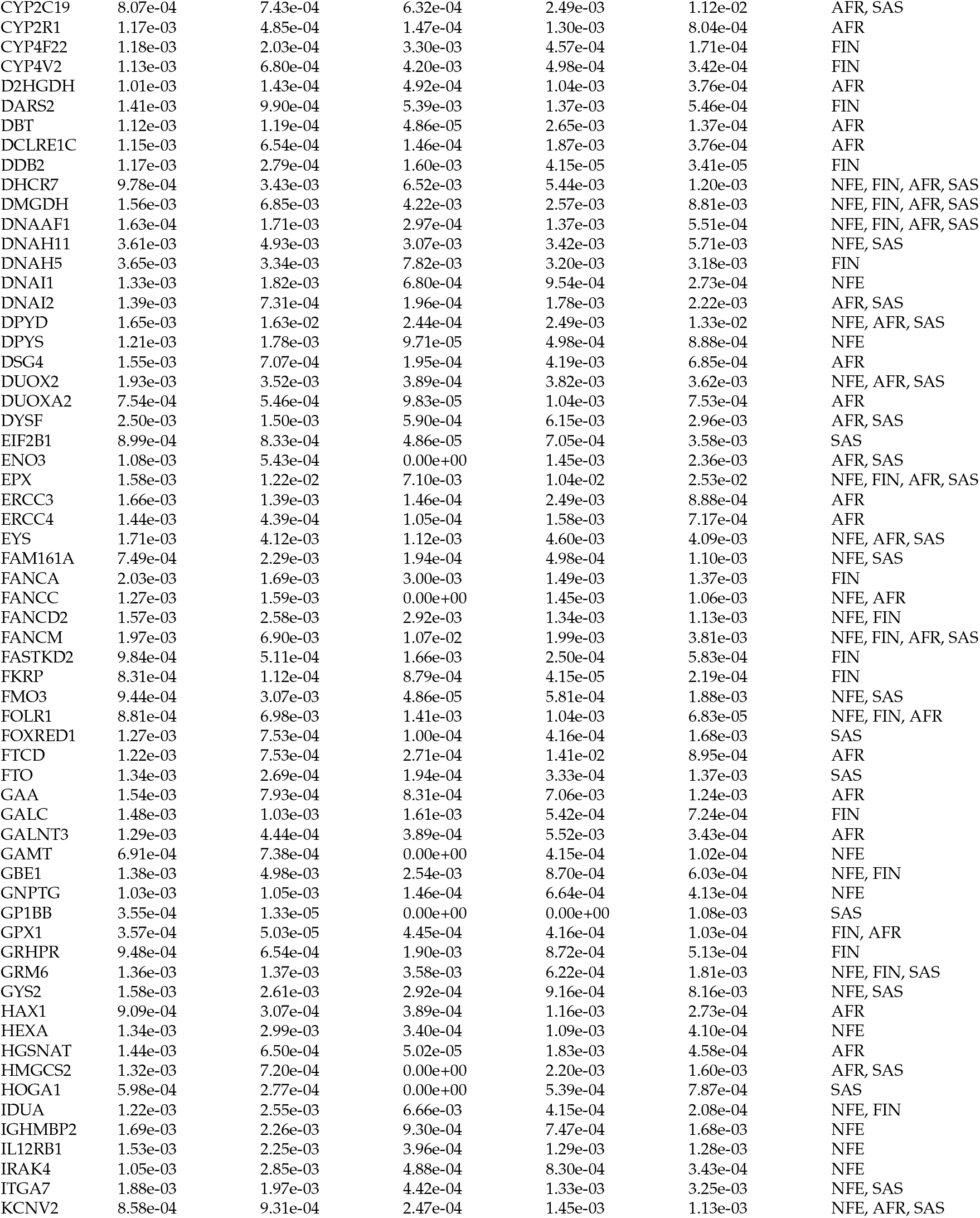
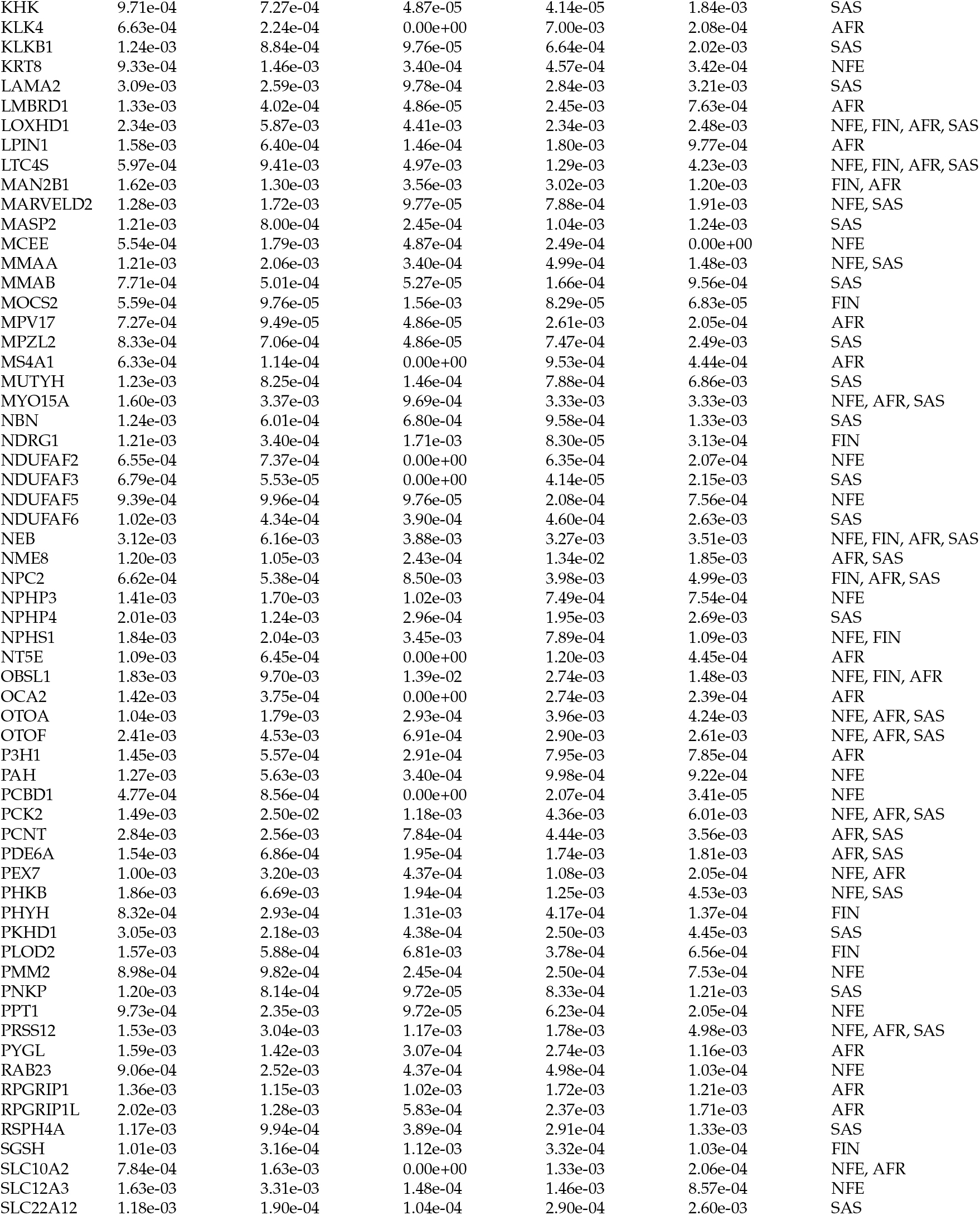
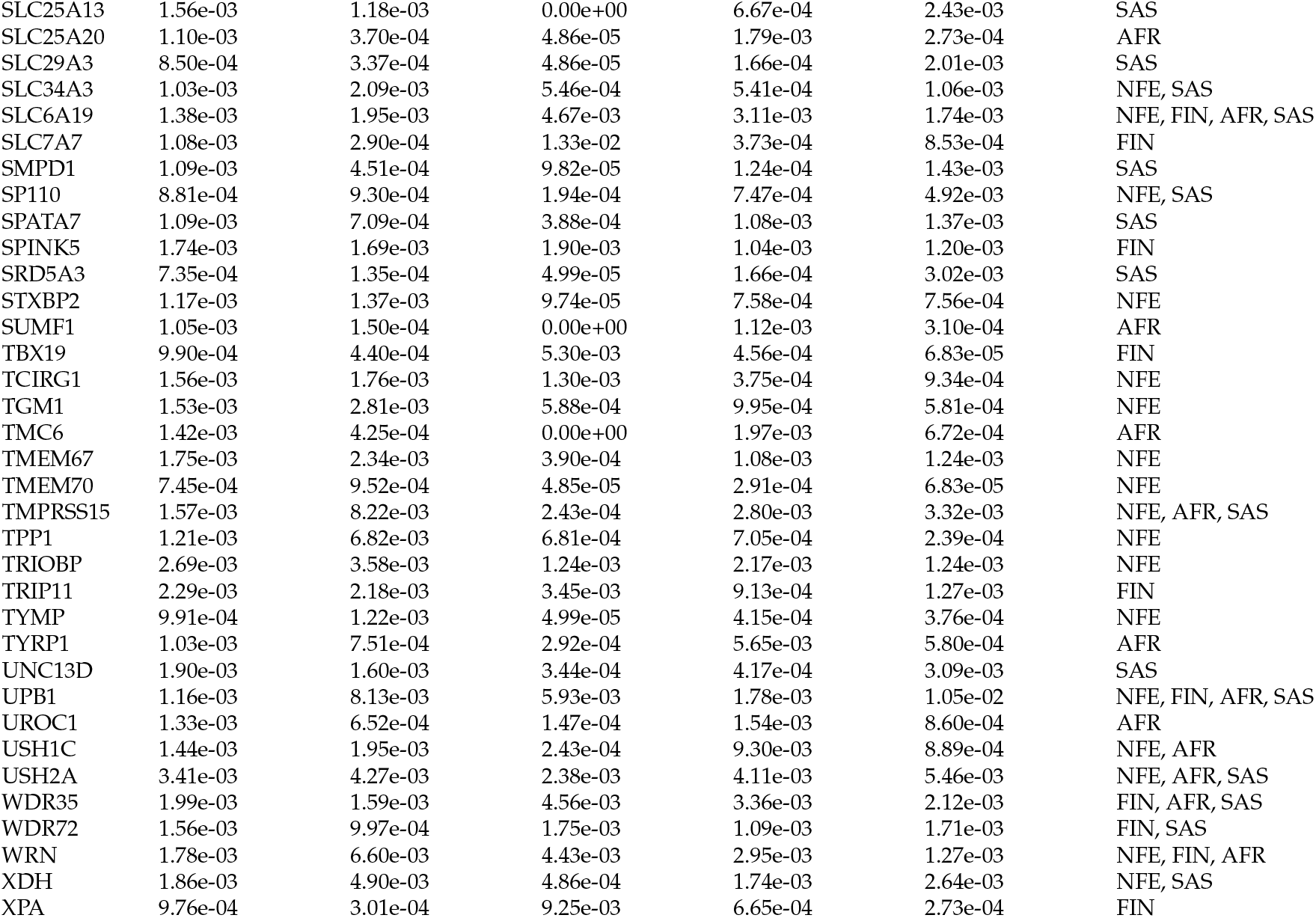
Potentially Purely Recessive Selection Genes Across Populations: List of recessive genes that appear to follow classic recessive mutation-selection balance based on at least one population-specific pLoF being larger than the gene’s square root mutational target size. Square root mutational target size, pLoFs for Non-Finnish European, Finnish, African, and South Asian populations, and populations where the gene matches mutation-selection balance model are included for each gene.

**Table S5:**
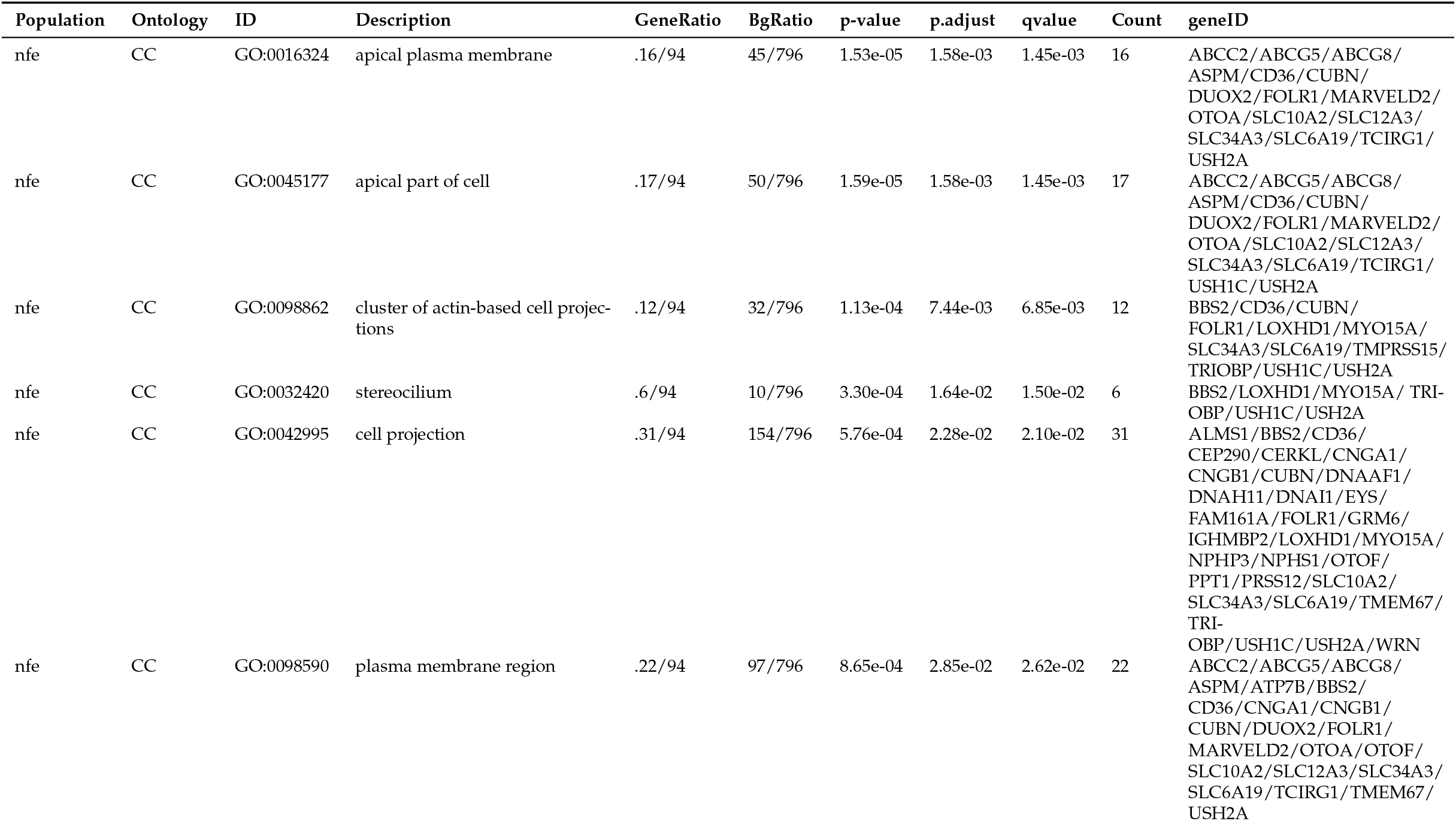
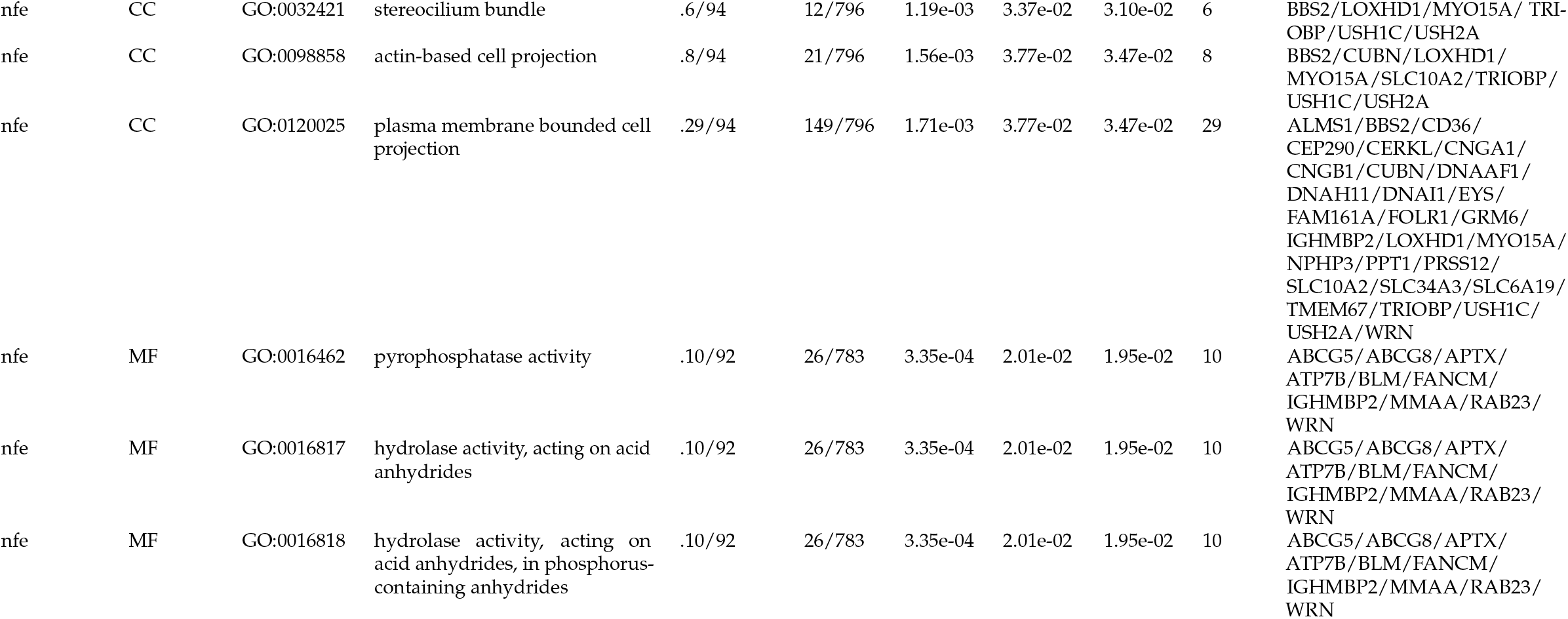
Select Recessive Genes GO Enrichment: Gene Ontology enrichment of recessive disease genes in the expected range for recessive mutation-selection balance based on pLoF in Non-Finnish European, Finnish, African, and South Asian populations. Each row identifies the enrichment of purely recessive genes in biological, molecular, and cellular functions and the significance of this enrichment. Background gene list is all recessive disease genes.

| Population | Ontology | ID | Description | GeneRatio | BgRatio | p-value | p.adjust | qvalue | Count | geneID |
| --- | --- | --- | --- | --- | --- | --- | --- | --- | --- | --- |
| nfe | CC | GO:0016324 | apical plasma membrane | .16/94 | 45/796 | 1.53e-05 | 1.58e-03 | 1.45e-03 | 16 | ABCC2/ABCG5/ABCG8/<br>ASPM/CD36/CUBN/<br>DUOX2/FOLR1/MARVELD2/<br>OTOA/SLC10A2/SLC12A3/<br>SLC34A3/SLC6A19/TCIRG1/<br>USH2A |
| nfe | CC | GO:0045177 | apical part of cell | .17/94 | 50/796 | 1.59e-05 | 1.58e-03 | 1.45e-03 | 17 | ABCC2/ABCG5/ABCG8/<br>ASPM/CD36/CUBN/<br>DUOX2/FOLR1/MARVELD2/<br>OTOA/SLC10A2/SLC12A3/<br>SLC34A3/SLC6A19/TCIRG1/<br>USH1C/USH2A |
| nfe | CC | GO:0098862 | cluster of actin-based cell projections | .12/94 | 32/796 | 1.13e-04 | 7.44e-03 | 6.85e-03 | 12 | BBS2/CD36/CUBN/<br>FOLR1/LOXHD1/MYO15A/<br>SLC34A3/SLC6A19/TMPRSS15/<br>TRIOBP/USH1C/USH2A |
| nfe | CC | GO:0032420 | stereocilium | .6/94 | 10/796 | 3.30e-04 | 1.64e-02 | 1.50e-02 | 6 | BBS2/LOXHD1/MYO15A/ TRI-OBP/USH1C/USH2A |
| nfe | CC | GO:0042995 | cell projection | .31/94 | 154/796 | 5.76e-04 | 2.28e-02 | 2.10e-02 | 31 | ALMS1/BBS2/CD36/<br>CEP290/CERKL/CNGA1/<br>CNGB1/CUBN/DNAAF1/<br>DNAH11/DNAI1/EYS/<br>FAM161A/FOLR1/GRM6/<br>IGHMBP2/LOXHD1/MYO15A/<br>NPHP3/NPHS1/OTOF/<br>PPT1/PRSS12/SLC10A2/<br>SLC34A3/SLC6A19/TMEM67/<br>TRI-OBP/USH1C/USH2A/WRN |
| nfe | CC | GO:0098590 | plasma membrane region | .22/94 | 97/796 | 8.65e-04 | 2.85e-02 | 2.62e-02 | 22 | ABCC2/ABCG5/ABCG8/<br>ASPM/ATP7B/BBS2/<br>CD36/CNGA1/CNGB1/<br>CUBN/DUOX2/FOLR1/<br>MARVELD2/OTOA/OTOF/<br>SLC10A2/SLC12A3/SLC34A3/<br>SLC6A19/TCIRG1/TMEM67/<br>USH2A |
| nfe | CC | GO:0032421 | stereocilium bundle | .6/94 | 12/796 | 1.19e-03 | 3.37e-02 | 3.10e-02 | 6 | BBS2/LOXHD1/MYO15A/ TRI-OBP/USH1C/USH2A |
| nfe | CC | GO:0098858 | actin-based cell projection | .8/94 | 21/796 | 1.56e-03 | 3.77e-02 | 3.47e-02 | 8 | BBS2/CUBN/LOXHD1/MYO15A/SLC10A2/TRIOBP/USH1C/USH2A |
| nfe | CC | GO:0120025 | plasma membrane bounded cell projection | .29/94 | 149/796 | 1.71e-03 | 3.77e-02 | 3.47e-02 | 29 | ALMS1/BBS2/CD36/CEP290/CERKL/CNGA1/CNGB1/CUBN/DNAAF1/DNAH11/DNAI1/EYS/FAM161A/FOLR1/GRM6/IGHMBP2/LOXHD1/MYO15A/NPHP3/PPT1/PRSS12/SLC10A2/SLC34A3/SLC6A19/TMEM67/TRIOBP/USH1C/USH2A/WRN |
| nfe | MF | GO:0016462 | pyrophosphatase activity | .10/92 | 26/783 | 3.35e-04 | 2.01e-02 | 1.95e-02 | 10 | ABCG5/ABCG8/APTX/ATP7B/BLM/FANCM/IGHMBP2/MMAA/RAB23/WRN |
| nfe | MF | GO:0016817 | hydrolase activity, acting on acid anhydrides | .10/92 | 26/783 | 3.35e-04 | 2.01e-02 | 1.95e-02 | 10 | ABCG5/ABCG8/APTX/ATP7B/BLM/FANCM/IGHMBP2/MMAA/RAB23/WRN |
| nfe | MF | GO:0016818 | hydrolase activity, acting on acid anhydrides, in phosphorus-containing anhydrides | .10/92 | 26/783 | 3.35e-04 | 2.01e-02 | 1.95e-02 | 10 | ABCG5/ABCG8/APTX/ATP7B/BLM/FANCM/IGHMBP2/MMAA/RAB23/WRN |

## Appendix

### Appendix A: Theoretical results

Here we derive our approximate formula for mutation-selection-drift balance in the purely recessive case in the presence of inbreeding. We also discuss how it generalizes previous results and discuss various computational considerations.

One of our main results in the course of deriving this theorem is rigorously showing that a discrete-time Wright-Fisher model with strength of selection *s*_*hom*_ against homozygotes and inbreeding controlled by a rate of identity-by-descent, *φ*, converges to the same Wright-Fisher diffusion as a the standard Wright-Fisher model *without inbreeding* but with an effective population size of *N*/(1 + *φ*), an effective strength of selection against *heterozygotes* of *φs*_*hom*_, and an effective strength of selection against homozygotes of (1 + *φ*)*s*_*hom*_.

#### A discrete-time Wright-Fisher model with inbreeding

We start by considering a single-locus biallelic discrete-time Wright-Fisher model. We fix the population size to be *N* diploids, and we label the two alleles “0” and “1”. Suppose that the frequency of the “1” allele in generation *t* is *f*_*t*_. The next generation is formed by first creating an infinitely large pool of gametes, where proportion *f*_*t*_ are the “1” allele. Then, each gamete independently mutates to the other allele with probability *µ*. We then consider pairing gametes to create zygotes. To model inbreeding, with probability *φ* a single gamete is pulled from the mutated gamete pool, and the two gametes in a zygote are identical to this chosen gamete. Otherwise, with probability (1 − *φ*) the two gametes are independently chosen from the mutated gamete pool. We then model purely recessive selection by removing zygotes homozygous for the “1” allele from the pool of zygotes with probability *s*_*hom*_. Finally, the next generation is formed by randomly choosing *N* of these zygotes from the pool. This process defines a Markov chain on the frequency of the “1” allele. The goal of this section is to approximately compute the mean of this frequency at equilibrium.

#### From the discrete-time Wright-Fisher model to a Wright-Fisher diffusion process

Our first step will be to move from this unwieldy discrete-time Wright-Fisher model to a more analytically tractable diffusion. Following [95, Chapter 15.1], to prove that this process converges to a diffusion, we need to compute 𝔼 [*f*_*t*_ − *f*_*t*−1_| *f*_*t*−1_] and 𝔼 [(*f*_*t*_ − *f*_*t*−1_)^2^| *f*_*t*−1_] and show that both are *O*(1/*N*), and we need to prove that 𝔼 [(*f*_*t*_ − *f*_*t*−1_)^4^| *f*_*t*−1_] is *o*(1/*N*). To calculate these quantities, we first introduce the notation 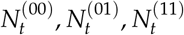 for the number of individuals in generation *t* who are homozygous for the “0” allele, heterozygous, and homozygous for the “1” allele respectively. Under our model, the number of “1” alleles, 2*N f*_*t*_ can be decomposed as 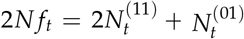.

Then, by construction we have that

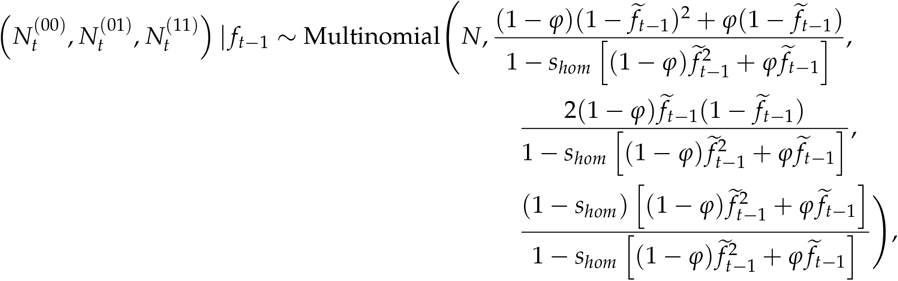

where

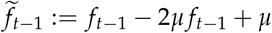

represents the frequency of the “1” allele in the gamete pool following mutation. Throughout we will assume — as is standard in population genetics — that *µ* and *s*_*hom*_ are *O*(1/*N*). Given that later we will apply our results to even the homozygous lethal case (*s*_*hom*_ = 1), we will revisit this assumption when numerically checking the accuracy of our approximations.

Therefore,

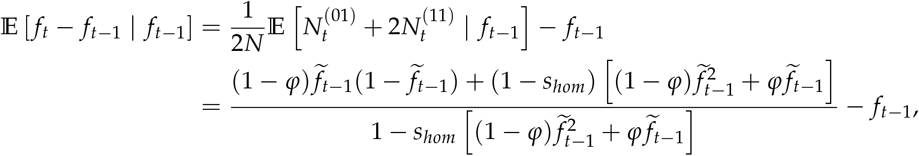

which after much algebra simplifies to

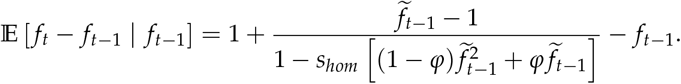

Using the series expansion 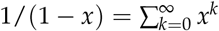, which is convergent for *x* ∈ (−1, 1), we obtain

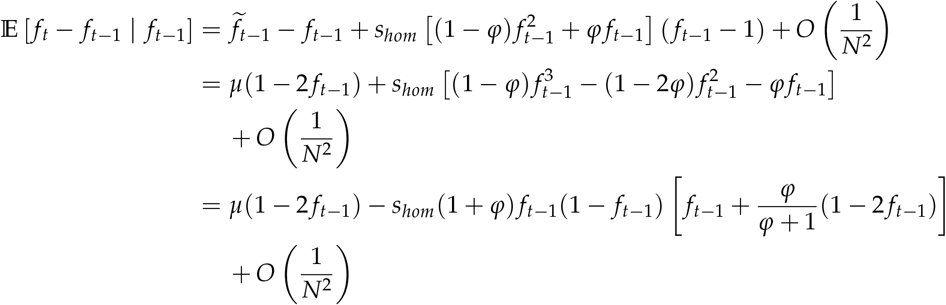

where in the first line we used that 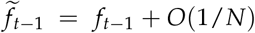. The slightly strange form of the final line shows that this expected change is equivalent (up to lower order terms) to a Wright-Fisher model with an *effective* strength of selection against heterozygotes, 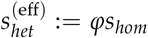, and an *effective* strength of selection against homozygotes of 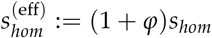. Overall, this is *O*(1/*N*) because of our assumptions on *µ* and *s*_*hom*_, and therefore the first moment of the change in allele frequency is consistent with convergence to a diffusion.

Similarly, using that for (*X*_1_, *X*_2_, …) ~ Multinomial(*N, p*_1_, *p*_2_, …),

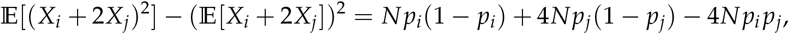

we can derive

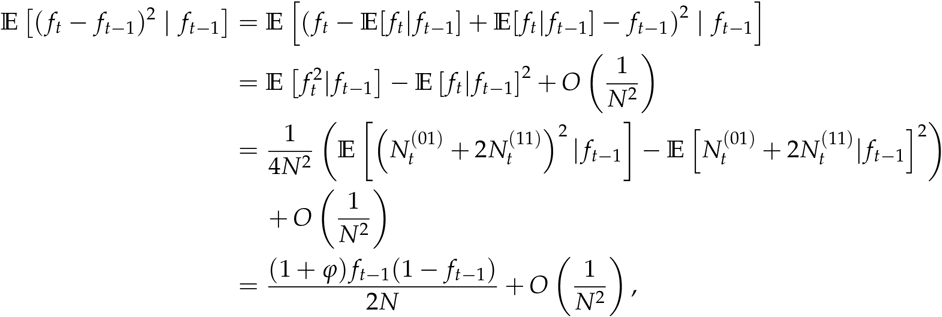

where the second equality relied on our result that the mean change in allele frequency is *O*(1/*N*), and to obtain the final equality we used similar algebra and arguments as in the case of the first moment. This is a factor of (1 + *φ*) larger than the second moment of the allele frequency change under a standard Wright-Fisher diffusion, indicating that genetic drift is (1 + *φ*) times stronger than in a population without inbreeding.

Finally, we can show that the fourth moment is small by considering:

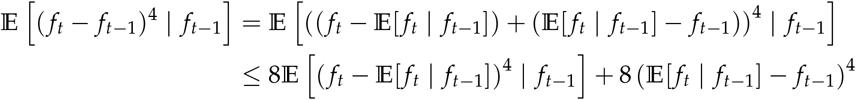

where the inequality can be derived from Young’s inequality or via convexity of *f* (*x*) : *x* ↦ *x*^4^. Continuing, we see

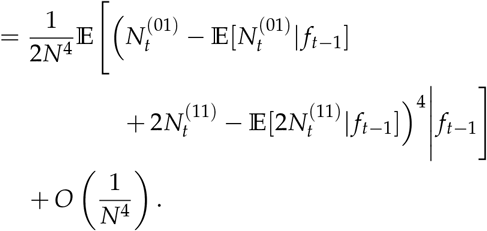

Applying Young’s inequality again, we obtain

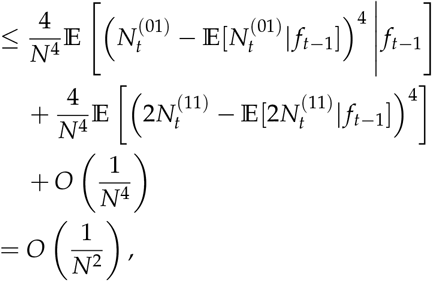

where the final line follows because the fourth central moment of a Binomial(*N, p*) random variable is *O*(*N*^2^). This shows that the fourth moment of the change in allele frequencies is *o*(1/*N*) as required to converge to a diffusion.

With these results in hand, we can now use the results of [95, Chapter 15.1] to show that in the large *N* limit this converges to a diffusion. In theory, this involves scaling time by a factor of *N* and keeping *Nμ* and *Ns*_*hom*_ fixed as *N* gets large. Here, we will (somewhat) abuse notation, by doing this scaling and taking the limit to obtain the diffusion and then immediately undoing the scaling so that we retain *µ* and *s*_*hom*_ in interpretable units. This results in the population frequency satisfying the following stochastic differential equation (SDE):

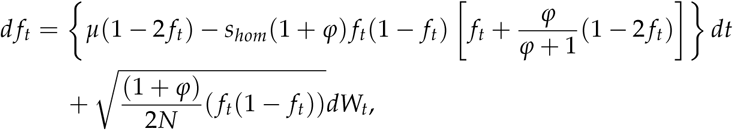

where *W*_*t*_ is a Wiener process (Brownian motion).

#### Stationary distribution of Wright-Fisher diffusion

The advantage of having obtained this SDE is that we may now readily derive the equilibrium probability density function. First, we can appeal to the Kolmogorov forward equation (Fokker-Plank equation) to transform the SDE into a partial differential equation for the probability density function, which we will denote by *p*(*f, t*). In particular, the Kolmogorov forward equation applied to our SDE implies

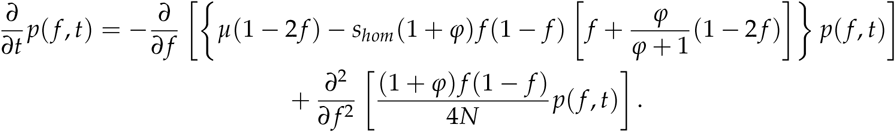

At equilibrium, there should be no change in the density with respect to time. We will denote the equilibrium density by *p*(*f*). Equating the PDE with zero and integrating with respect to *f* results in

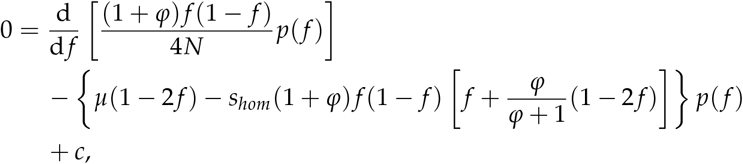

for some integration constant *c*, which can be shown to be zero by an argument about the “flux” of probability. Essentially, one can interpret the right hand side of the previous equation without the constant as the amount of probability mass “flowing through” the point *f*. Since the process is at equilibrium and exists on a bounded domain, and mass cannot “flow” past the boundaries, this implies that *c* = 0. Using this and that d/d *f* [*p*(*f*)]/*p*(*f*) = d/d *f* log *p*(*f*), and rearranging we arrive at

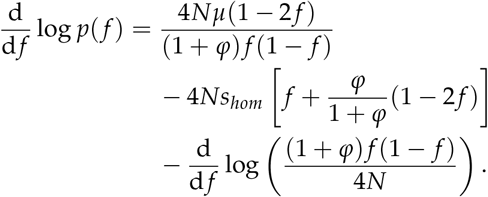

This can now be readily solved up to the normalizing constant by direct integration, resulting in

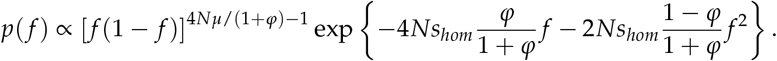

#### A simplification under the assumption of strong selection

Unfortunately, obtaining the normalizing constant for this density is intractable. To make progress, we assume that *Ns*_*hom*_ is large so that *f* is small with very high probability. In this case, we drop the (1 −*f*) term, and instead of assuming that *f*∈ [0, 1], we instead allow any *f* ≥ 0, with the understanding that the probability that *f >* 1 is vanishingly small. This is intimately related to previous work and has connections to branching processes [96, 97]. In any case, ultimately we obtain

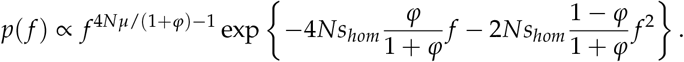

#### Main analytical results and numerical considerations

Below, in Theorem 1 we prove a more general result that may be of independent interest that implies that at equilibrium the expected frequency under our simplified model is

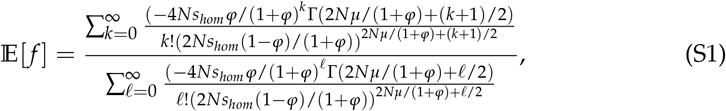

where Γ(·) is the Gamma function.

In the limit of no inbreeding, this can be easily seen to reduce to

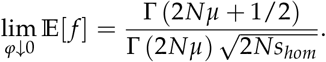

Computing 𝔼 [*f*] under our model with *φ* = 0 was one of the main results of [96], showing that our results are consistent with and generalize those.

The series appearing in the numerator and denominator of Equation S1 are convergent, and so this formula is “exact” (up to the diffusion and small *f* approximations needed to derive the density function in the first place). Unfortunately, the terms in the series can be astronomically large and have alternating signs. As a result, in our implementation of Equation S1, we use `mpmath` [98], a high precision library in python, and we compute all of the Gamma functions and factorials in log-space and perform cancellations before exponentiating. Furthermore, we typically truncate the series after a number of terms. We adaptively truncate the series by computing terms until the next term in the series is smaller in magnitude than the previous term and smaller than exp(™500).

Another downside of Equation S1 is that it provides little intuition into how the various parameters affect the equilibrium frequency without evaluating the equation. As a result, we also prove a general result, Theorem 2, about the large *N* limit. As we will show below, when *φ* is small, *N* must be quite large for this to be a good approximation (e.g., *N* ≫ 10,000). In our case, this result implies that

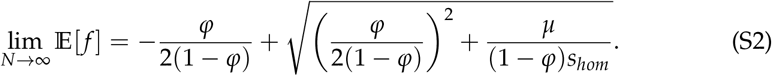

This equation is much easier to reason about directly. For example, we see that when *φ* is 0, this reduces to 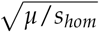, the classic mutation selection balance result used in the main text. Similarly, when *φ* is large, this is well approximated by *µ*/*s*_*hom*_, as can be seen via Taylor series. This is intuitive, because this would be the result of mutation selection balance if selection were acting on *heterozygotes* with strength *s*_*hom*_. In essence, when inbreeding is strong, almost all individuals are homozygotes, and so we can envision the population as being essentially *N* haploids, and selection acts with strength *s*_*hom*_ to remove haploids with the deleterious allele. Mutations can occur on either of an individual’s chromosomes, but both chromosome are equally likely to be passed on to their likely homozygous offspring. These factors of two cancel, resulting in the effective rate of mutation in this haploid model remaining *µ*.

#### Estimation of fraction of selection in heterozygotes

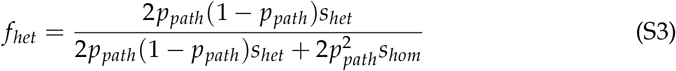

For 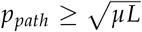 we set 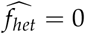. Otherwise, after some basic cancellations from the expression above, we estimate 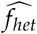 under the conservative assumption that *s*_*hom*_ = 1, as follows:

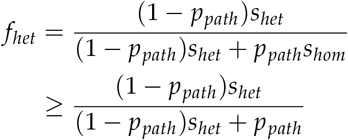

Noting that under mutation selection balance 2*p*(1 − *p*)*s*_*het*_ = 2*μL* 2*p*^2^*s*_*hom*_, and again setting *s*_*hom*_ = 1, we rearrange the latter expression to get:

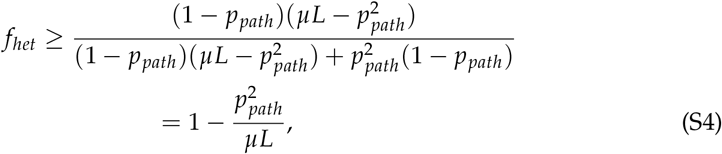

We use the latter as a lower-bound estimator for *f*_*het*_.

#### Verification of approximations using fastDTWF

In addition to the comparisons to simulations presented in the main text, we also checked Equation S1 and Equation S2 against the equilibrium values obtained numerically using `fastDTWF` [99]. `fastDTWF` uses numerical approximations to compute the probability mass function of having different allele frequencies under geneal discrete-time Wright-Fisher models, including the one described at the beginning of this appendix. It uses principled numerical approximations, but does not perform any of the approximations to the model that we performed to obtain our main results (e.g., taking the diffusion limit or assuming that the frequencies are small with high probability).

The results are shown in Figures A1 to A3 where we vary *N, µ, s*_*hom*_, and *φ*. Unless otherwise stated, we use a default value of *µ* = 10^−5^, *s*_*hom*_ = 0.1, and *φ* = 0.02. Overall, it is clear that as long as the product of *N* and *s*_*hom*_ is large enough, Equation S1 is extremely accurate even for fairly small *N* and large *s*_*hom*_, where the assumptions we made during our derivation break down. For example, the results for *s*_*hom*_ = 1 are shockingly accurate given that our derivation required *s*_*hom*_ = *O*(1/*N*).

**Figure A1:**
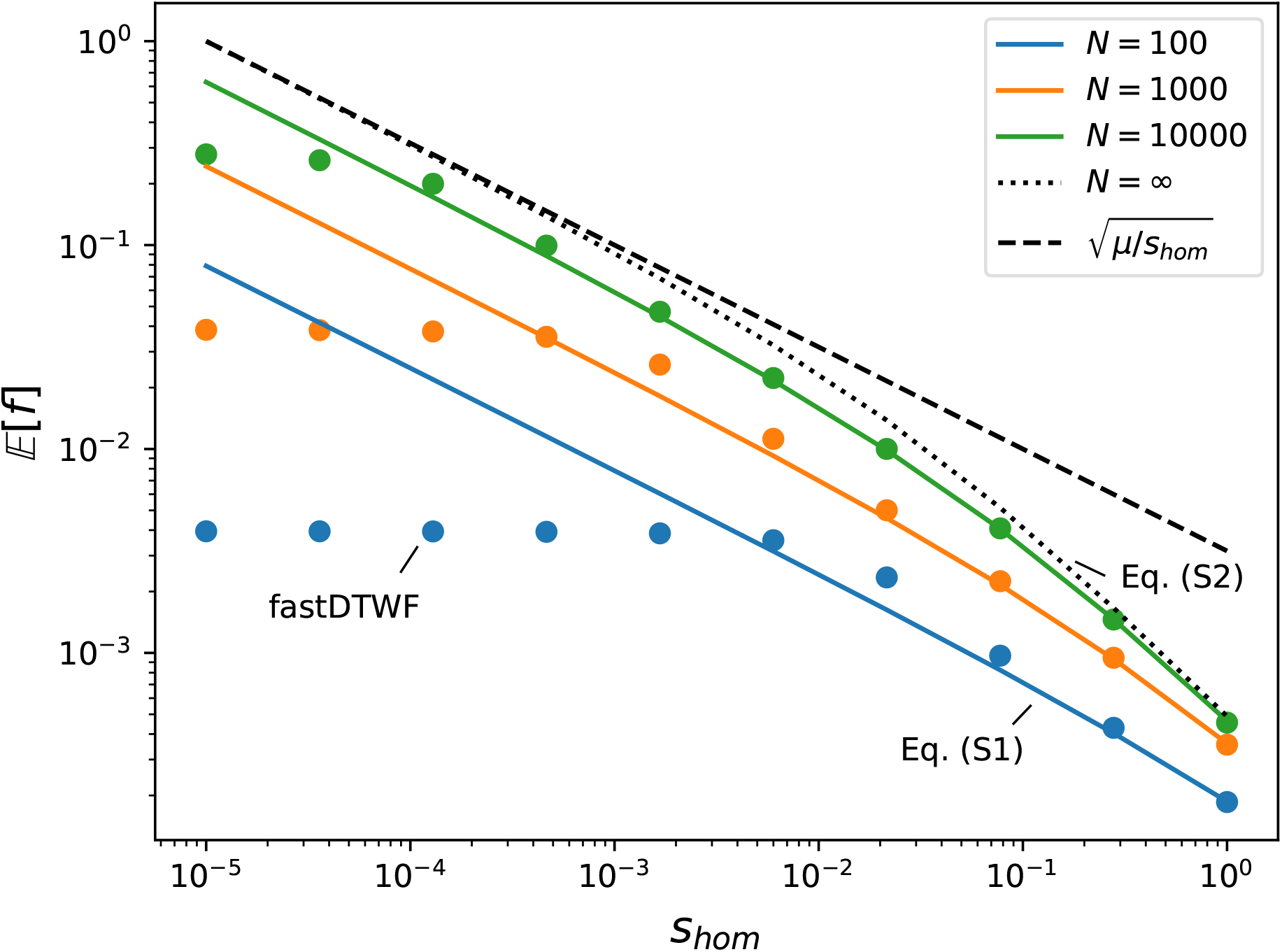
Comparison of `fastDTWF` with Equations S1 and S2, varying *N* and *s*_*hom*_. The other parameters of the model are fixed at *µ* = 10^−5^ and *φ* = 0.02. `fastDTWF` is presented as dots and the results derived in this appendix are lines. The colored solid lines are from Equation S1, the dotted line is the *N* → ∞ asymptotic result from Equation S2, and the dashed line is the asymptotic mutation-selection balance result without inbreeding, 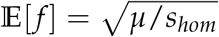.

**Figure A2:**
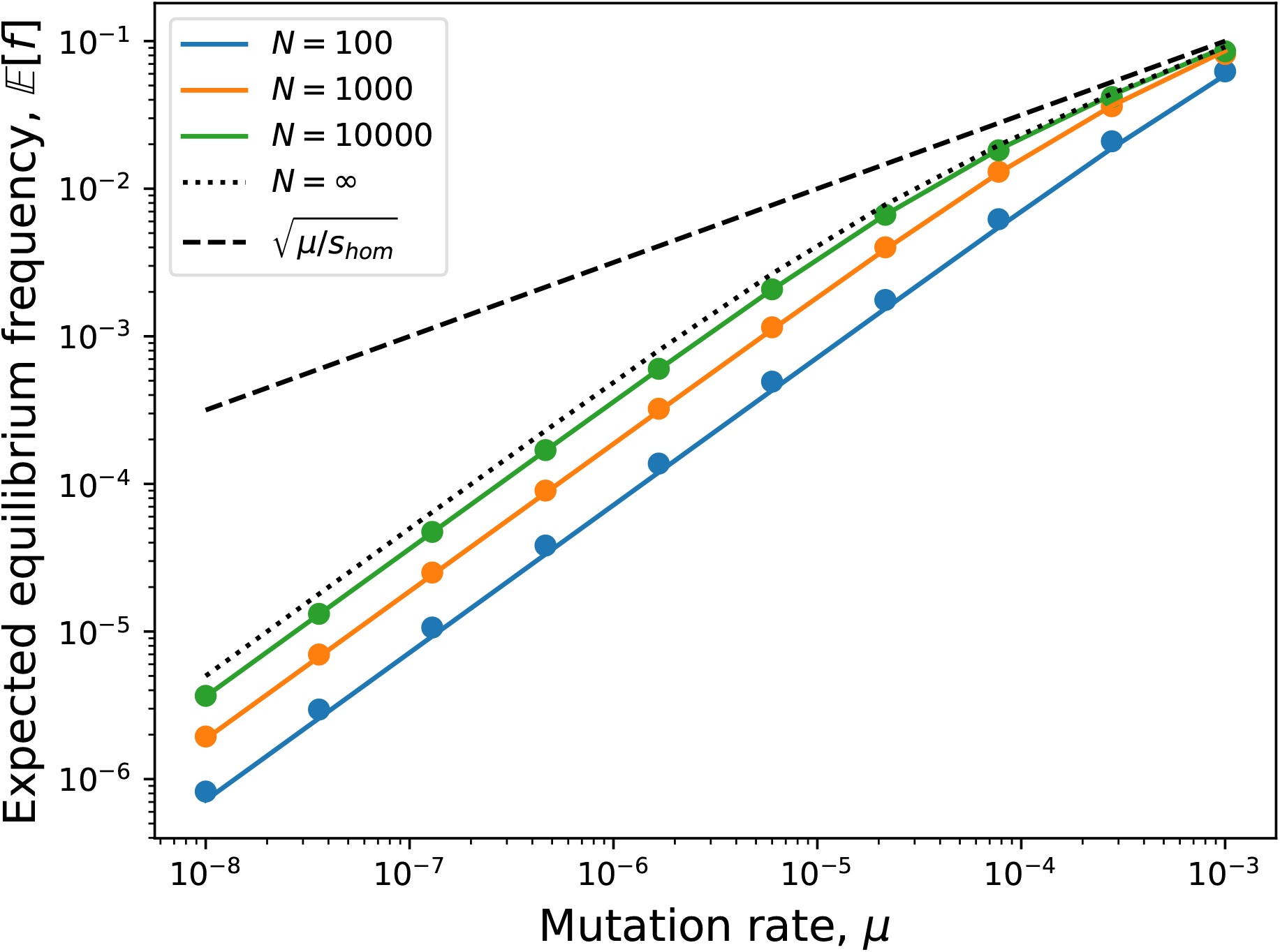
Comparison of `fastDTWF` with Equations S1 and S2, varying *N* and *µ*. The other parameters of the model are fixed at *s*_*hom*_ = 0.1 and *φ* = 0.02. `fastDTWF` is presented as dots and the results derived in this appendix are lines. The colored solid lines are from Equation S1, the dotted line is the *N* → ∞ asymptotic result from Equation S2, and the dashed line is the asymptotic mutation-selection balance result without inbreeding, 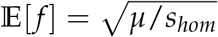.

**Figure A3:**
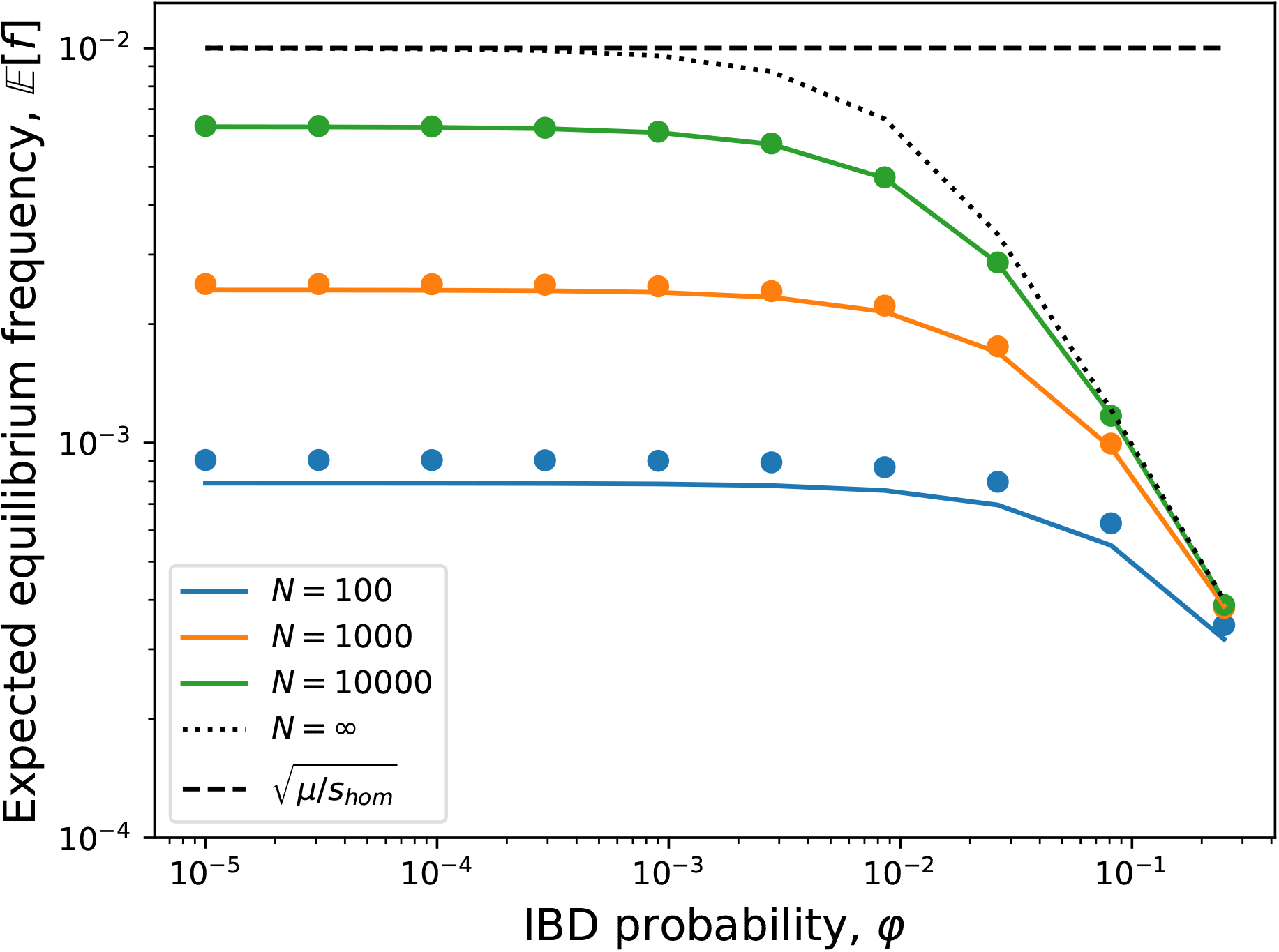
Comparison of `fastDTWF` with Equations S1 and S2, varying *N* and *φ*. The other parameters of the model are fixed at *s*_*hom*_ = 0.1 and *µ* = 10^−5^. `fastDTWF` is presented as dots and the results derived in this appendix are lines. The colored solid lines are from Equation S1, the dotted line is the *N* → ∞ asymptotic result from Equation S2, and the dashed line is the asymptotic mutation-selection balance result without inbreeding, 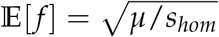.

#### Technical results

Here we prove our two technical results. The first theorem is fairly general and may be of independent interest, while the second Theorem is an easy application of Laplace’s method to our setting.

##### Theorem 1.

*Let X be a random variable supported on* [0, ∞) *drawn from a distribution with probability density function*

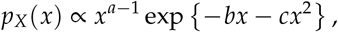

*for some a >* 0, *b >* 0 *and c >* 0. *Then*,

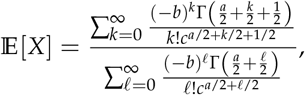

*where* Γ(·) *is the gamma function*.

*Proof*. To begin, we note that

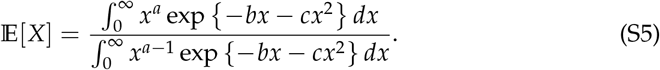

We will now obtain explicit formulas for the numerator and denominator. The approach for both is identical, so here we just derive the numerator. Substituting the series expansion of exp {−*bx*} into the numerator, we obtain

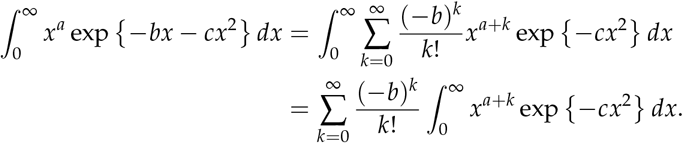

Changing the order of integration and summation on the second line requires justification, which we will provide at the end of this proof. At this point, we perform the change of variables *y* = *cx*^2^ to obtain

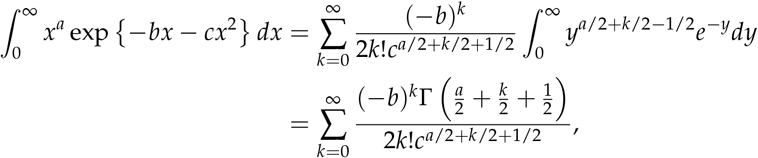

where the final line follows by the integral definition of the gamma function. We can apply exactly the same logic to the denominator of the right hand side of Equation S5, substituting *a ™*1 for *a*. Plugging the resulting series into Equation S5 and canceling factors of 2 proves the formula in the theorem.

To formally justify swapping the order of summations above, we appeal to Fubini’s Theorem, which can be applied if the series is absolutely convergent. This follows from noting

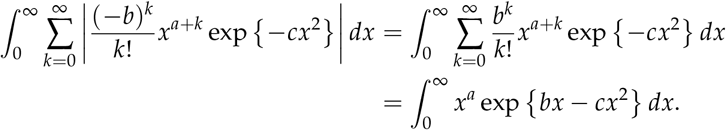

To bound this integral, we note that *bx ™cx*^2^ ≤−*bx* + *b*^2^/*c* by concavity and considering the tangent at *x* = *b*/*c*. Therefore,

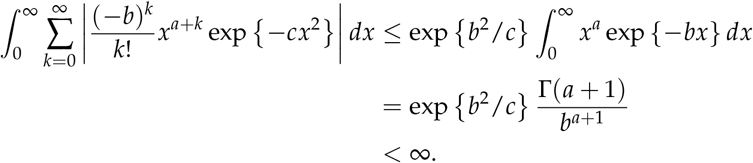

This also proves that the series appearing in the equation in the theorem are convergent. Additionally, the denominator is strictly positive, as can be seen by noting that the integrands in Equation S5 are strictly positive. As a result the series expression for 𝔼 [*X*] is well-defined. □

##### Theorem 2.

*Let X be a random variable supported on* [0, ∞) *drawn from a distribution with probability density function*

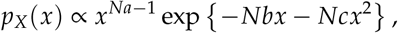

*for some a >* 0, *b >* 0 *and c >* 0. *Then, asymptotically as N* → ∞,

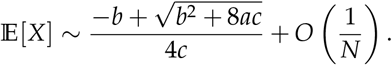

*Proof*. As in Theorem 1, we begin by noting that

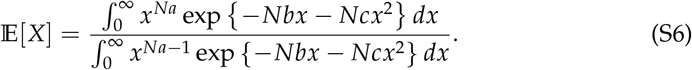

The proof is then a simple application of Laplace’s method (e.g., [100, Chapter 6.4]) to the numerator and denominator of Equation S6, which proceeds as follows. To handle the numerator and denominator simultaneously, we will consider the exponent of *x* in the integrands to be *Na ™δ* where *δ* = 0 in the numerator and 1 in the denominator. We then rewrite the integral as

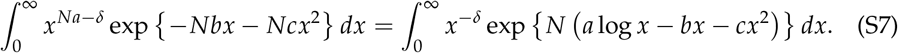

Consider the function *ψ*(*x*) := *a* log *x* − *bx* − *cx*^2^. Computing the first two derivatives,

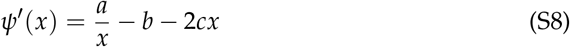

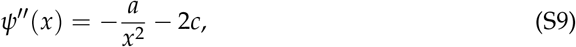

we see by Equation S9 that *ψ* is concave on the relevant domain of *x*, and by setting Equation S8 to zero and solving, we see that *ψ* has a global maximum over *x >* 0 at 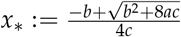.

Now, as *N* → ∞ values of *x* away from *x*_*_ contribute exponentially less to the integral in Equation S7. We may therefore truncate the limits of integration to a small, but fixed region around *x*_*_ (say of width 2*ϵ* for some tiny *ϵ* that is constant in *N*) without changing the asymptotic value of the integral:

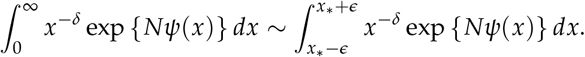

This in turn, allows us to Taylor expand *x*^−*δ*^ and *ψ*(*x*) about *x*_*_:

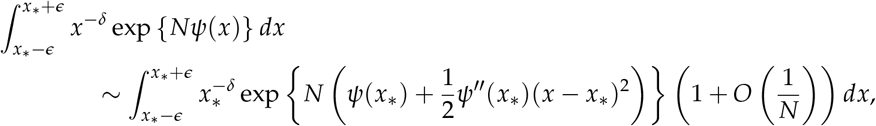

where the first order term *ψ*′(*x*_*_)(*x ™x*_*_) was dropped because it is zero as *x*_*_ is the maximizer of *ψ*(*x*). We will explain why the error term is *O*(1/*N*) at the end of the proof. Next, we again use the fact that the contributions to this integral outside of our window of width 2*ϵ* are asymptotically negligible, but this time we use this observation to expand our window of integration:

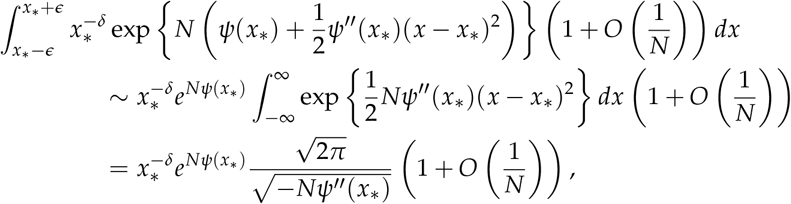

where the final line follows from recognizing the integrand as the unnormalized density of Gaussian random variable with mean *x*_*_ and variance ™1/*ψ*′′(*x*_*_). The proof follows from substituting this expression with *δ* = 0 and *δ* = 1 into the numerator and denominator of Equation S6 respectively.

We now explain why the error term is *O*(1/*N*). Further terms in the Taylor series for *ψ*(*x*) can be collected into exp{*N*(*ψ*′′′(*x*_*_)(*x* − *x*_*_)^3^ + · · ·)}, which can then itself be expanded as 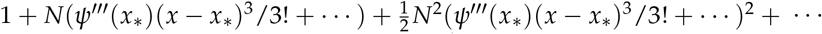. Multiplying this series with the Taylor series of *x*^−*δ*^ about *x*_*_ results in a series in powers of (*x* − *x*_*_). We will call the coefficient of (*x* − *x*_*_)^*k*^ in this expansion *A*_*k*_, where we can see that 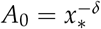. Furthermore, *A*_2_ is *O*(1), *A*_4_ is *O*(*N*) and for *k* ≥ 6, *A*_*k*_ is at most *O*(*N*^*k*/3^). We then have

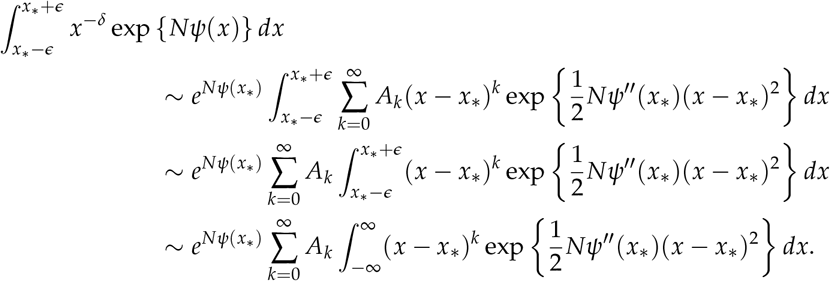

We then have that all odd order terms vanish because they are proportional to odd central moments of a Gaussian random variable. Similarly, the even terms are proportional to even central moments of a Gaussian with variance *O*(1/*N*), and hence the integral involving (*x* − *x*_*_)^2*k*^ is *O*(1/*N*^*k*^). That is,

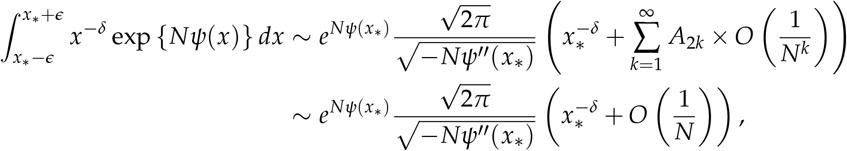

where the final line follows from noting that the terms involving *A*_2_ and *A*_4_ are ≤ *O*(1/*N*) and all other terms are strictly lower order by our bound that *A*_2*k*_ ≤ *O*(*N*^2*k*/3^) for *k* ≥ 3, and hence *A*_2*k*_ × *O*(1/*N*^*k*^) ≤ *O*(*N*^−*k*/3^) ≤ *O*(1/*N*) for *k* ≥ 3. □

